# GPCR endocytosis rewires neuronal gene expression and cellular architecture

**DOI:** 10.1101/2025.08.26.672159

**Authors:** Katherine L. Hall, Matthew J. Klauer, Malarie E. Schexnider, Nikoleta G. Tsvetanova

## Abstract

In the brain, G protein-coupled receptors (GPCRs) regulate neuronal excitability, synaptic transmission, and behavior by engaging transcriptional and translational programs that produce enduring changes in cellular function and architecture. However, the molecular mechanisms that couple GPCR activation to these adaptations remain poorly understood. Here, we demonstrate that the beta-adrenergic receptor (β2AR), a mediator of noradrenaline in the central nervous system, remodels neuronal morphology through compartmentalized signaling pathways that orchestrate distinct layers of gene regulation. Following stimulation, β2ARs remain active on endosomes, and their intracellular signaling promotes dendritic growth and synapse formation in human iPSC-derived and primary cortical neurons. These structural effects are driven by two coordinated regulatory axes: PKA/CREB-dependent transcription of morphogenesis-related genes and PKA/mTOR-dependent translation of components of the protein synthesis machinery. Altogether, this work defines novel spatial and biochemical principles by which GPCR signaling drives structural reorganization and functional adaptations in neurons.

## INTRODUCTION

G protein-coupled receptors (GPCRs) make up the largest and most diverse family of membrane receptors in humans. The vast majority of non-sensory GPCRs are expressed in the brain where they regulate neuronal excitability and synaptic plasticity to support key functions, including learning, memory, reward processing, and motor control^1^. Not surprising, disruptions in GPCR signaling have been linked to a wide range of neurological and psychiatric disorders, making these receptors major targets for neuropharmacological therapies^1–3^. Despite their widespread relevance, how GPCRs drive long-term changes in neuronal function remains poorly defined. These adaptations depend on precise regulation of gene expression, including transcriptional and translational reprogramming that reshapes neuronal identity and connectivity. Such regulatory events are particularly important for sustained forms of plasticity, such as long-term potentiation^4,5^. Yet, the nature of these gene expression programs, and how these are coordinated downstream of GPCRs, have not been systematically characterized.

At the cellular level, GPCRs transduce their effects by activation of heterotrimeric G proteins and regulation of the abundance of second messengers. The cyclic AMP-protein kinase A (cAMP-PKA) cascade is a major signaling pathway engaged by neuronal GPCRs. Upon ligand binding, receptors coupled to the G protein, Gαs, stimulate adenylyl cyclase, leading to increase in intracellular cAMP. Elevated levels of cAMP in turn activate PKA and other effectors, which influence a broad range of cellular targets and neuronal processes, including development and excitability^6^. Achieving this functional diversity requires mechanisms that can direct signals with both spatial and temporal precision. Recent studies suggest that the subcellular location of GPCR activation contributes to this specificity. In particular, GPCR signaling is not restricted to the plasma membrane, but can continue from intracellular compartments, prolonging and/or shaping activation of select downstream pathways^7–10^. Although endosomal signaling has been described for several neuronal GPCRs, much of the mechanistic understanding of compartmentalized Gαs/cAMP signaling derives from heterologous or non-neuronal systems^9^. Therefore, whether these principles govern long-term gene regulation and cellular responses in neurons has remained unclear.

Given the size and morphological complexity of neurons, we reasoned that compartmentalized Gαs/cAMP signaling may be particularly important for relaying extracellular signals across extended cellular domains and into specialized compartments to regulate gene expression. We addressed this question using the beta2-adrenergic receptor (β2AR), a prototypical Gαs-coupled GPCR and well-established model for localized cAMP signaling. The β2AR is one of the most extensively studied GPCRs and is expressed across multiple tissues, including the brain. In the central nervous system, it mediates key noradrenergic effects on synaptic plasticity by engaging transcription- and translation-dependent mechanisms that regulate attention, arousal, learning, and memory^11–14^. In addition, prior work in HEK293 cells demonstrated that β2AR remains active following endocytosis^15,16^, making it an ideal system to determine whether endosomal Gαs/cAMP signaling is conserved in neurons, and how it contributes to long-term neuronal remodeling. Here, we integrate imaging-based analyses of dendritic and synaptic architecture with global transcriptional and translational profiling to show that β2AR activation induces extensive structural remodeling through compartment-specific regulation of gene expression.

## RESULTS

### **β**2AR signaling promotes dendritogenesis and synapse formation

Because cortical excitatory neurons are a primary site of adrenergic receptor function^14,17–19^, we employed a previously validated human iPSC-derived neuronal system (iNeurons) that expresses β2AR but no other β-adrenergic receptor subtypes (**Fig. 1a, Supplementary Fig. 1a)**^20^. These neurons are generated by neurogenin 2-driven differentiation and acquire a cortical glutamatergic identity^21^. We chose iNeurons because they provide a unique human neuronal platform for dissecting GPCR signaling and downstream gene regulation. In addition to biological relevance, the scalability and homogeneity of iNeurons make them well-suited for robust transcriptomic and translatomic profiling.

**Figure 1.**
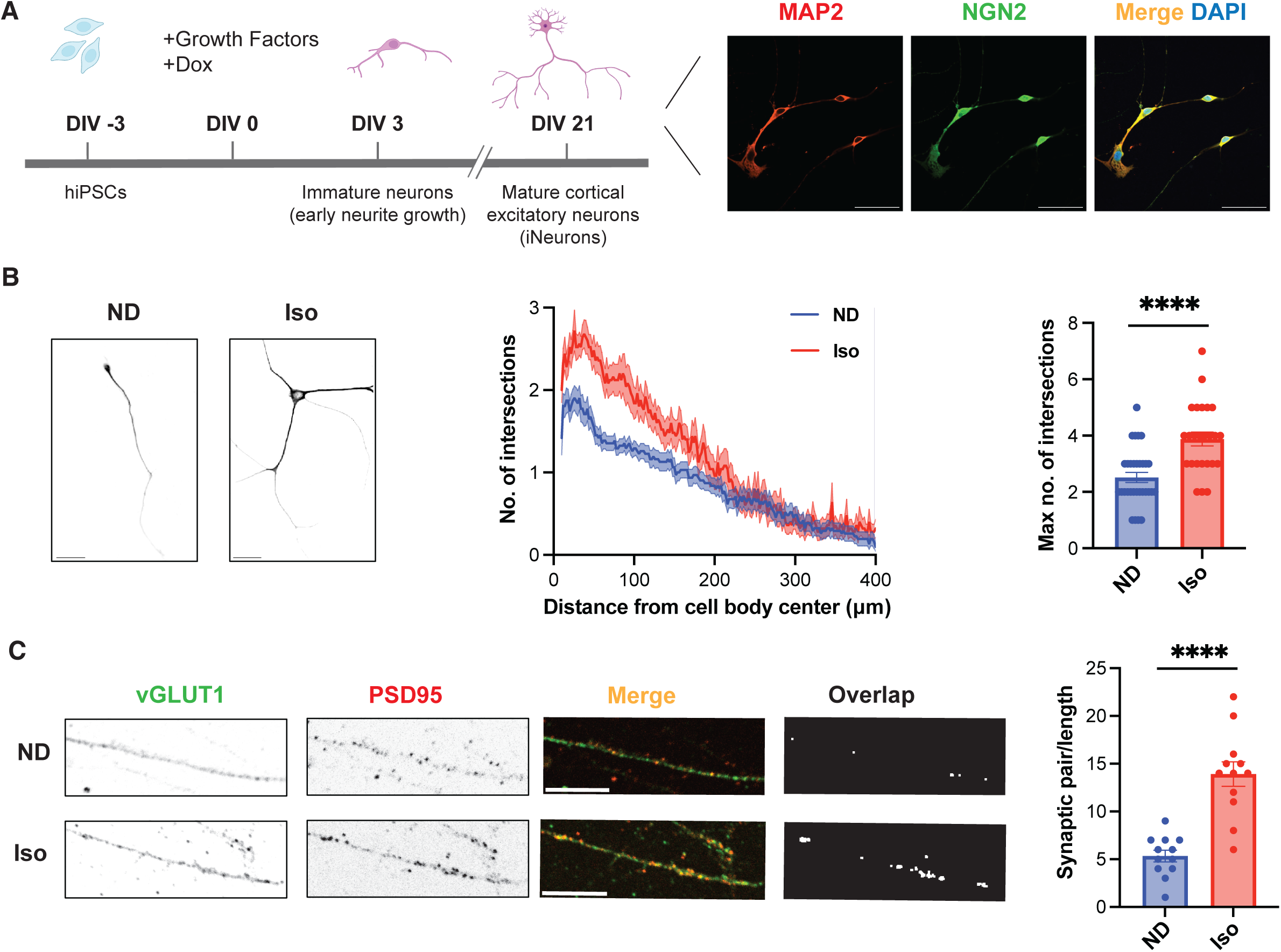
β2AR activation promotes structural remodeling of neurons. **(A)** *Left*: Schematic of the workflow for generation of human iPSC-derived cortical neurons. *Right*: Fixed cell fluorescence microscopy staining of iNeuron cultures (DIV21) with antibodies for the cortical marker, NGN2 (green), and neuronal marker, MAP2 (red). **(B)** Representative images of neurons treated with vehicle (no drug, ‘ND’) or 1 μM Isoproterenol (Iso) for 2 h, fixed and stained with antibody for MAP2 three days later *(left)*; quantification of maximum number of crossings by Sholl analysis *(right)*. Data are mean of n = 21-25 cells from 3 independent experiments. **(C)** *Left:* Representative images of neurons stained for vGLUT1 as presynaptic marker (green) and PSD95 as postsynaptic marker (red). *Right:* Quantification of synaptic pairs based on colocalization analysis of vGLUT1/PSD95 puncta per 35 μ*m*. Neurons were treated with vehicle (‘ND’) or 1 μM Iso for 24 h. Data are mean of n = 12 cells from 3 independent experiments. Error bars represent standard error of the mean. **** = *p* < 0.0001 by unpaired Student’s *t* test. ND = no drug. Scale bars, 50 μm in **(B)** and 10 μm in **(C)**.

β2AR activity promotes excitability and synaptic potentiation, processes that are generally coupled to structural remodeling of the neuron. We therefore first asked whether β2AR signaling has effects on dendritic growth and synapse formation by stimulating iNeurons with the synthetic beta-adrenergic receptor agonist, isoproterenol. To quantify the effects of β2AR signaling on dendritic development, we focused on an early stage of neuronal differentiation, when individual dendritic arbors can be accurately reconstructed and quantified. We performed Sholl analysis which revealed a significant ligand-dependent increase in the number of dendritic branches (**Fig. 1b, Supplementary Fig. 1b)**. To measure synapse formation, we quantified colocalization of the pre- and postsynaptic markers, VGLUT1 and PSD95, respectively. Treatment of iNeurons differentiated for 3 weeks with isoproterenol caused an increase in the number of synaptic pairs (**Fig. 1c, Supplementary Fig. 1c)**. We observed similar effects on neuronal morphogenesis following treatment with the selective β2AR agonist salmeterol and the endogenous catecholamine norepinephrine **(Supplementary Fig. 1d-e)**. Therefore, activation of the β2AR enhances both dendritic complexity and synapse formation, indicating a prominent role of the cascade in promoting neuronal structural remodeling. Next, we sought to define the spatial organization of the β2AR signaling pathway and the molecular programs that underlie its role in neuronal morphogenesis.

### Intracellular **β**2AR signaling is required for neuronal structural remodeling

It was previously reported in HEK293 cells that the β2AR initiates G protein-dependent signaling from the plasma membrane and from endosomes^15,16^. We first examined receptor trafficking in iNeurons and found that, internalized β2ARs colocalized with the early endosomal marker EEA1 following isoproterenol stimulation **(Supplementary Fig. 2a)**. To assess the activity of β2ARs in neuronal compartments, we used two validated single-chain antibody (nanobody) sensors, Nb80 and Nb37, that recognize the active state of the receptor and Gαs protein, respectively^15^. Each biosensor tagged with green fluorescent protein (GFP) was co-expressed with the β2AR in iNeurons. Under unstimulated conditions, the receptor was confined to the plasma membrane, while the nanobody was diffusely distributed in the cytoplasm by live confocal microscopy (**Fig. 2a-b)**. Following stimulation with isoproterenol, both biosensors co-localized with the internalized receptor on endosomal vesicles (**Fig. 2a-b)**. These support that β2AR and Gαs are active on endosomes in neurons. To provide evidence that intracellular activation contributes to the resulting cellular response, we measured receptor-induced cAMP production. To estimate the contribution of endosomal signaling to the overall cellular cAMP response, we restricted β2AR signaling to the plasma membrane by acute endocytic inhibition with the compound Dyngo-4a, an established method to block dynamin-dependent trafficking **(Supplementary Fig. 2b)**^15,16^. Blockade of receptor endocytosis significantly blunted agonist-induced cAMP accumulation (**Fig. 2c)**. Hence, signaling by internalized β2ARs contributes to the cumulative cellular cAMP response.

**Figure 2.**
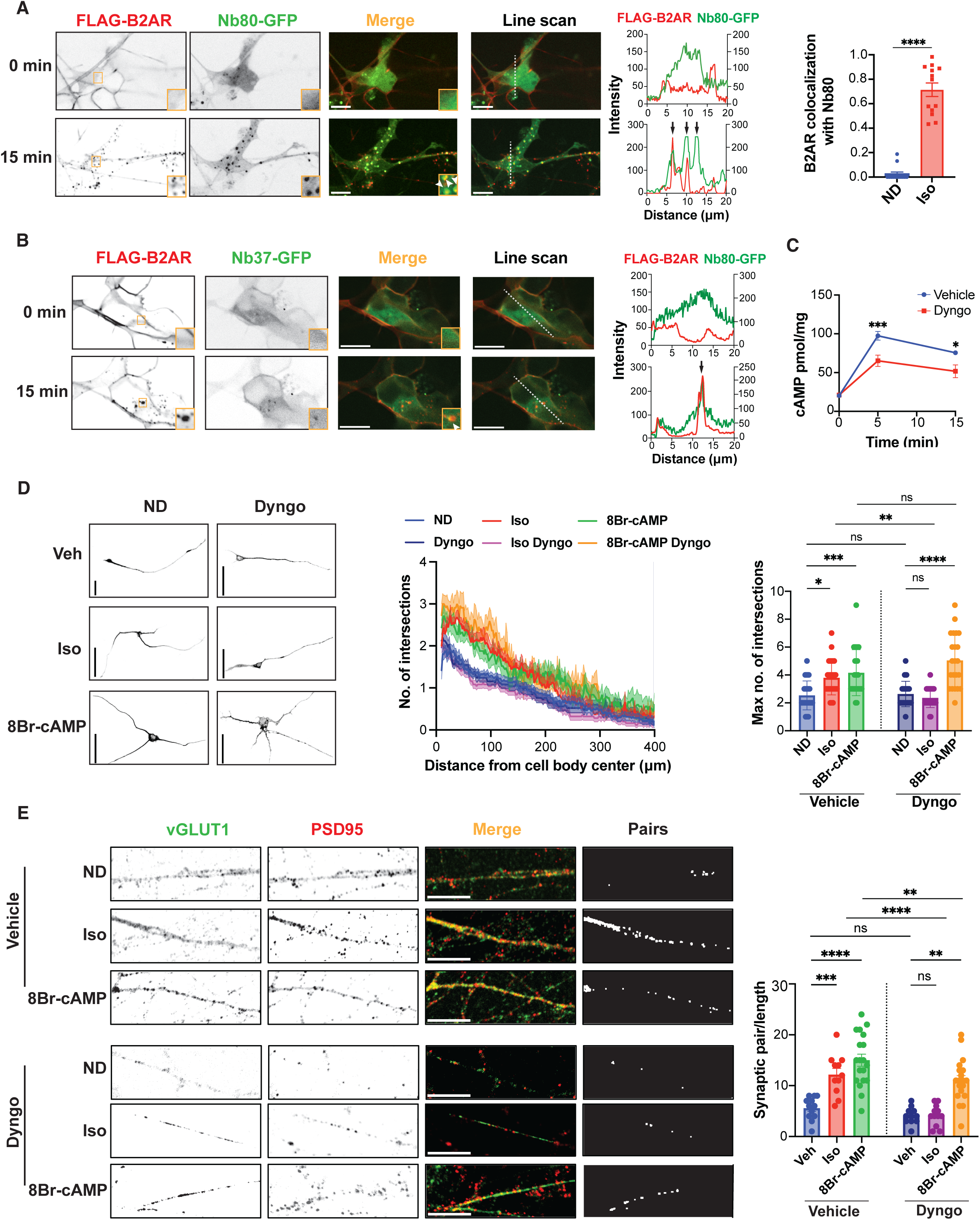
Endosomal signaling by β2AR is required for dendrite growth and synapse formation. **(A)** *Left*: Representative confocal images of neurons expressing Flag-tagged receptor (red) and GFP-tagged conformational biosensors for detection of active β2AR (Nb80, green); 1 μM Isoproterenol (Iso) was added at t=0. *Middle*: Line scan analyses of colocalization between the β2AR and the biosensor. White dotted line: line scan analysis through the center of the cell of gray value in each channel. Arrows indicate colocalization. *Right:* Mander’s coefficient calculated for β2AR colocalization with Nb80 plotted between conditions, n =14 cells in 3 independent experiments. **(B)** *Left*: Representative confocal images of neurons expressing Flag-tagged receptor (red) and GFP-tagged conformational biosensors for detection of active Gαs (Nb37, green); 1 μM Isoproterenol (Iso) was added at t=0. This is a representative image of n =10 cells in 3 independent experiments. *Right*: Line scan analyses of colocalization between the β2AR and the biosensor. White dotted line: line scan analysis through the center of the cell of gray value in each channel. Arrows indicate colocalization. **(C)** Endocytosis is required for cAMP accumulation. Neurons were pretreated with vehicle (DMSO) or 30 μM Dyngo-4a for 30 min, then stimulated with 1 μM Iso for 5 min or 15 min, lysed and cAMP levels were measured with an ELISA assay. Data are mean of n = 3 independent experiments. **(D-E)** Analyses of dendritogenesis and synapse formation. Representative images are shown for each condition. Neurons were fixed and stained for MAP2 **(D)** or vGLUT1/PSD95 **(E)**. Neurons were pretreated with vehicle (DMSO) or 30 μM Dyngo-4a for 30 min, then stimulated with 1 μM Iso or 100 μM cAMP analog (8-Br-cAMP). Quantification of number of crossings by Sholl analysis **(D)** and synaptic pairs **(E)** is shown. Data are mean of n = 18-25 cells in **(D)** and n = 11-19 cells in **(E)** from 3 independent experiments. Error bars represent standard error of the mean. **** = *p* < 0.0001, *** = *p* < 0.001, ** = *p* < 0.01, * = *p* < 0.05 by unpaired Student’s *t* test **(A)**, two-way ANOVA with Sidak correction in **(C-E)**. ND = no drug; Veh = vehicle. Scale bars, 10 μm in **(A), (B), (E)** and 50 μm in **(D)**.

Next, we asked whether the intracellular β2AR population is necessary to induce dendrite growth and synapse formation. Remarkably, pre-treatment of cells with Dyngo-4a completely abolished the effects of isoproterenol on neuron morphology (**Fig. 2d-e)**. Notably, direct activation of the cAMP pathway with a cell-permeable cAMP analog, 8-Br-cAMP, retained a robust morphogenic effect even with endocytic blockade (**Fig. 2d-e)**. Because pharmacological inhibition of dynamin disrupts cellular trafficking more broadly and can have pleiotropic effects, we sought an orthogonal approach to selectively interfere with endosomal β2AR signaling. We adapted a previously established genetic strategy that relies on endosomal targeting of Nb80 to restrict β2AR activity^22^. Because Nb80 binds the same intracellular receptor interface engaged by Gαs, it can serve not only as a biosensor for receptor activity, but also as a steric inhibitor of receptor-G protein coupling when concentrated at a specific membrane compartment. We therefore targeted Nb80 to endosomes by fusion to tandem FYVE domains (2xFYVE), which bind PI3P-enriched endosomal membranes, and used nuclear-targeted Nb80 (NLS-Nb80) as a spatial control **(Supplementary Fig. 2c)**. Expression of endosome-targeted Nb80 abolished isoproterenol-induced dendritic growth and synapse formation, whereas NLS-Nb80 had no effect **(Supplementary Fig. 2d-e)**. Together, these complementary approaches demonstrate that signaling by the endosomal β2AR population is required for neuronal structural remodeling.

### **β**2AR activation induces extensive neuronal transcriptional and translational reprogramming

Changes in neuronal morphology depend on de novo protein synthesis to provide the molecular machinery needed for structural adaptation. Notably, these processes are regulated by both transcription-dependent and transcription-independent mechanisms, because proteins required for synapse remodeling can be translated from both newly transcribed and pre-existing mRNA^23^. Therefore, to determine the molecular underpinnings of β2AR-dependent structural remodeling, we set out to delineate the global neuronal transcriptional and translational programs regulated by the receptor.

To identify the β2AR-dependent transcriptional responses, we conducted RNA sequencing (RNA-seq) analysis of iNeurons treated with isoproterenol for 2 hours. Through differential expression analysis, we identified ∼400 neuronal transcriptional targets, the majority of which were upregulated by agonist (**Fig. 3a, Dataset S1)**. Comparison with our previous analyses in non-neuronal cell models^16,24^ revealed that only a small subset of these genes had been identified as β2AR transcriptional targets (39/402 genes, *p* < 2.0 x 10^-16^ by Fisher’s exact test) and likely represent a conserved receptor-dependent transcriptional program **(Dataset S1)**. These included well-established cAMP-responsive immediate early genes (IEGs) such as *FOS*, *CGA*, and *DUSP1* **(Supplementary Fig. 3a)**. In contrast, more than 90% of the β2AR-regulated transcriptome was unique to iNeurons, revealing a substantially broader and highly cell type-specific transcriptional response than previously appreciated. This neuron-specific program encompassed numerous established regulators of neurite outgrowth, synaptic development, and neuronal activity, including *BDNF, NPAS4, NEUROG2, BTBD3, SYT4, NPTX1, NPTX2, ELFN1, UNCX,* and *KCNA2*. Consistent with this, Gene Ontology analysis identified enrichment for biological processes associated with neuronal development, including “neuron differentiation”, “neuropeptide signaling”, and “anatomical structure development” (**Fig. 3b)**. This strong enrichment for neuronal developmental pathways was consistent with the profound effects of β2AR signaling on neuronal morphogenesis (**Fig. 1, Supplementary Fig. 1)**. Among these targets, we selected *BDNF*, a key regulator of axon growth and guidance, dendritic arborization, and synaptic connectivity^25^, and *UNCX*, a transcription factor implicated in neuronal differentiation and connectivity^26^, for experimental validation. Agonist-dependent induction of both *BDNF* and *UNCX* was confirmed by RT-qPCR **(Supplementary Fig. 3a)**. These results demonstrate that, although β2AR signaling engages a conserved core transcriptional program across cell types, the majority of receptor-regulated genes in neurons are tailored to the unique physiological functions of these cells.

**Figure 3.**
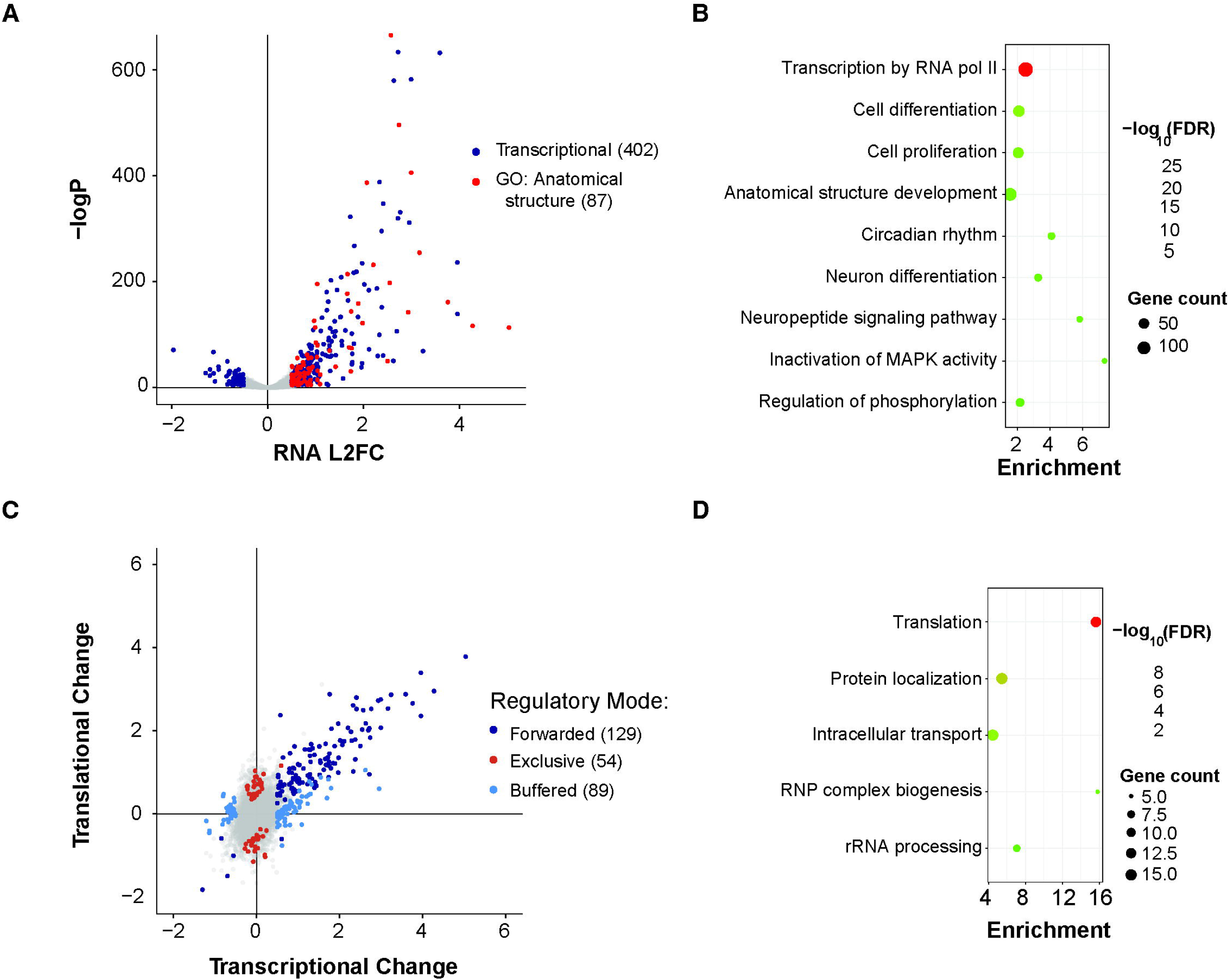
Transcriptional and translational responses to GPCR activation. **(A)** Volcano plot of RNA-seq analysis showing neuronal gene fold-induction following activation with 1 μM Isoproterenol for 2 h (log_2_ Isoproterenol/no drug). Blue dots represent β2AR-dependent transcriptional targets. Red dots represent transcriptional targets that make up the Gene Ontology category “Anatomical structure development” in **(B)**. Data are mean of n = 3 independent experiments. **(B)** Gene ontology categories enriched among the transcriptional targets. **(C)** Scatter plot of changes in mRNA (‘Transcriptional change’) and RPF abundance (‘Translational change’) following activation with 1 μM Isoproterenol for 2 h from integrated RNA-seq and Ribo-seq analyses. Values are log_2_ (Isoproterenol/no drug). Dark blue dots represent forwarded targets. Light blue dots represent buffered targets. Red dots represent exclusively translationally regulated targets. The parentheses indicate number of target genes. Data are mean of n = 3 (RNA-seq) or mean of n = 4 (Ribo-seq) independent experiments. **(D)** Gene ontology categories enriched among the exclusive translationally regulated β2AR-dependent targets. **(B)** and **(D)**: Color of the circle represents significance (-Log_10_FDR). Circle size represents number of gene counts. Plotted by SRplot^58^.

Next, we assessed how activation of the receptor cascade impacts the neuronal translational landscape. To this end, we carried out ribosome profiling (Ribo-seq) under matched stimulation conditions **(Supplementary Fig. 3b)**. The sequenced ribosome footprints (RPFs) exhibited the expected size distribution, three-nucleotide periodicity and enrichment primarily in coding sequences confirming that the analysis measured ribosome occupancy on actively translating neuronal mRNAs **(Supplementary Fig. 3c-d)**. To evaluate the extent of overlap between transcriptional and translational responses to β2AR, we first examined the correlation between RNA-seq and Ribo-seq replicates. Although RNA abundance and ribosome occupancy were broadly correlated, replicates clustered more closely within each assay than across assays, consistent with both shared and distinct layers of transcriptional and translational regulation **(Supplementary Fig. 3e**, compare scatter plots on top vs bottom).

Considering genes with uniquely mapped reads across the two experimental platforms, we compared agonist-dependent changes in RPF and RNA abundance to determine how β2AR signaling coordinated transcriptional and translational responses. Integration of the RNA-seq and Ribo-seq datasets revealed three distinct modes of gene regulation (**Fig. 3c)**. First, we identified 129 “forwarded” targets, for which agonist-induced changes in mRNA abundance were accompanied by proportional changes in ribosome occupancy (**Fig. 3c, dark blue; Dataset S2)**. These genes therefore underwent coordinated transcription and translation and are expected to produce corresponding changes in protein abundance. Notably, this group included numerous regulators of neuronal activity, differentiation and structural remodeling **(Dataset S3)**. In contrast, 89 genes exhibited significant changes in mRNA abundance without corresponding changes in ribosome occupancy (“buffered” targets; **Fig. 3c, light blue, Datasets S2)**. For these genes, β2AR-dependent changes in transcript abundance were not propagated to protein synthesis. Lastly, 54 genes exhibited altered ribosome occupancy in the absence of detectable changes in mRNA abundance and were classified as “exclusive” translational targets of β2AR (**Fig. 3c, red; Dataset S2)**. These mRNAs therefore underwent selective translational regulation and would not have been identified by transcriptomic analysis alone. Remarkably, one-third of these exclusively translated targets encoded components of the protein synthesis machinery (**Fig. 3d, Dataset S3)**. Representative genes from each regulatory class were validated by RT-qPCR and immunoblotting, confirming the distinct modes of regulation predicted by the integrated RNA-seq/Ribo-seq analysis **(Supplementary Fig. 3f-g)**.

Together, these genomic analyses demonstrate that β2AR signaling remodels neuronal gene expression through coordinated transcriptional and translational regulatory mechanisms. This multilayered regulation promotes the expression of genes involved in neuronal growth and connectivity while independently enhancing the translation of mRNAs encoding components of the protein synthesis machinery.

### GPCR endocytosis is required for coordinated gene expression control

Given that β2AR endocytosis was required for isoproterenol-driven morphogenesis, we hypothesized that the agonist-induced gene expression changes may be critically dependent on intracellular receptor signaling. To interrogate the contribution of endosomal β2ARs to differential gene expression, we conducted parallel transcriptional and translational analyses in unstimulated and isoproterenol-treated iNeurons, in which receptor trafficking was blocked acutely with Dyngo-4a. Application of inhibitor alone did not interfere with steady-state neuronal gene expression **(Supplementary Fig. 4a-b)**. By contrast, Dyngo-4a markedly suppressed the agonist-driven responses at both the transcriptional and translational levels (*p* < 2.0 × 10^-16^ for β2AR transcriptional targets; *p* < 7.0 × 10^-7^ for exclusive β2AR translational targets, Wilcoxon test).

To further understand the scope and nature of the regulation, we classified genes within each dataset as targets of endosomal or plasma membrane signaling based on inhibitor effects. This revealed that endocytosis was broadly required for β2AR-mediated transcriptional responses, as all targets underwent significant loss of regulation in the presence of Dyngo-4a (FDR < 0.1 by multiple *t-*test analysis, **Fig. 4a**). RT-qPCR analysis of select IEGs and the morphogenesis factor *UNCX* confirmed that their expression was regulated by β2AR in an endocytosis-dependent manner **(Supplementary Fig. 4c-d)**. We similarly stratified the exclusive translational hits according to the impact of Dyngo-4a on their translation efficiency. As with the transcriptional responses, endosomal β2ARs emerged as the primary drivers of translation and were required for the regulation of ∼85% of target RNAs, (**Fig. 4b**, “endo”). These included genes that fell into the significant Ribo-seq GO categories identified in cells with intact endocytosis, including “translation” **(Supplementary Fig. 4e)**. However, while none of the transcriptional targets exhibited intact regulation upon endocytic blockade, we found that the regulation of ∼15% of translational targets did not require internalization. Hence, these RNAs represented targets of plasma membrane β2ARs (**Fig. 4b**, “PM”).

**Figure 4.**
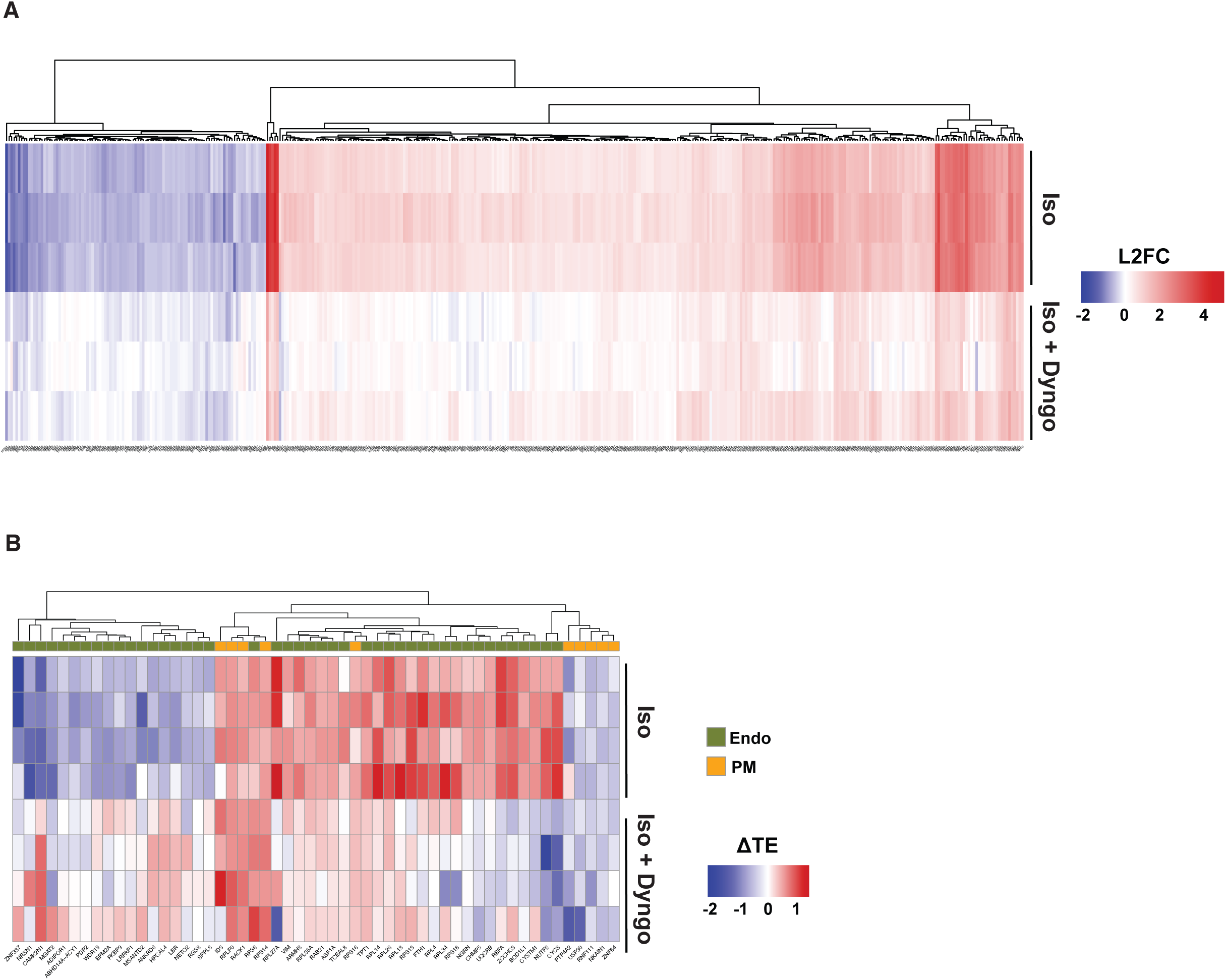
β2AR-induced gene reprogramming requires endocytosis. **(A-B)** Heatmaps of transcriptional **(A)** and exclusive translational **(B)** targets from RNA-seq/Ribo-seq analyses. Neurons were pretreated with vehicle (DMSO) or 30 μM Dyngo-4a for 30 min, then stimulated with 1 μM Isoproterenol (Iso) for 2 h. Colors represent log_2_ fold changes between Iso-treated and the corresponding untreated samples in terms of mRNA abundance **(A)** or changes in translation efficiency (ΔTE, the ratio between Iso-dependent changes in ribosomal occupancy and transcript abundance). Color-coding in **(B)** indicates whether a gene is classified as endosomal (‘Endo’, green) or plasma membrane target (‘PM’, yellow) based on multiple unpaired *t* test analysis with FDR < 0.1 cut-off. K-means clustering analysis was performed with pheatmap in R.

Together, these findings indicated that both transcriptional and translational responses to β2AR activation are predominantly endosome-dependent, although the translational regulation of a subset of mRNAs persisted even in the absence of intracellular receptors.

### Endosomal **β**2ARs remodel neuronal morphology through PKA/CREB and PKA/mTOR signaling axes

To uncover which molecular pathways are engaged by endosomal receptor activity to drive gene regulation, we analyzed motif enrichment patterns across target gene sets. Transcription factor analysis^27,28^ revealed CREB as the major driver of β2AR-induced transcriptional signaling **(Supplementary Fig. 5a)**. Pharmacological inhibition of CREB with compound 666-15^29^ abolished isoproterenol-induced transcriptional responses and prevented agonist-dependent dendritic growth and synapse formation (**Fig. 5a-b**, green bars; **Supplementary Fig. 5b)**, demonstrating that CREB mediates the morphogenic program downstream of β2AR signaling. Because CREB is canonically activated by PKA^16^, we next examined whether endosomal β2AR signaling engages this pathway. Using genetically encoded biosensors targeted to the cytosol and nucleus^30,31^, we found that isoproterenol robustly activated PKA in both compartments and that this response was significantly blunted by Dyngo-4a **(Supplementary Fig. 5c-d)**. Correspondingly, isoproterenol increased phosphorylation of CREB on Ser133, an effect abolished by inhibition of either PKA or receptor endocytosis **(Supplementary Fig. 5e)**. Moreover, pharmacological inhibition of PKA prevented receptor-dependent transcriptional induction of *UNCX* and IEGs **(Supplementary Fig. 5f-g)**. Ultimately, inhibition of PKA abolished isoproterenol-induced dendritic growth and synapse formation (**Fig. 5a-b, red bars)**, demonstrating that PKA is the primary effector of structural adaptations elicited by β2ARs.

**Figure 5.**
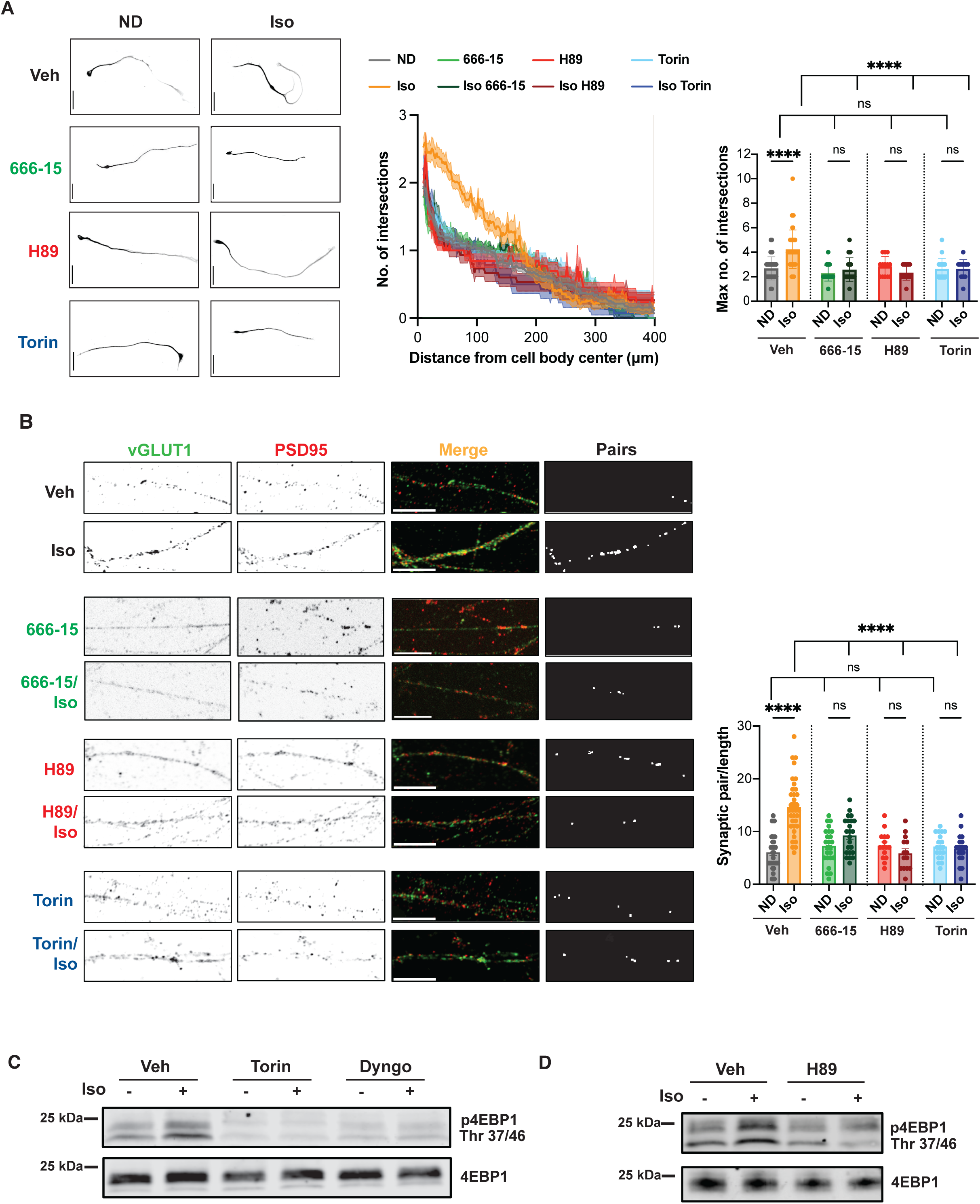
Compartmentalized β2AR activation modulates neuronal architecture via distinct PKA-linked pathways. **(A-B)** Analyses of dendritogenesis and synapse formation under various treatment conditions. Neurons were pretreated with vehicle (DMSO), 1 μM 666-15, 10 μM H89 or 100 nM Torin for 30 min, then stimulated with 1 μM Isoproterenol (Iso). Neurons were fixed and stained for MAP2 **(A)** or vGLUT1/PSD95 **(B)**. *Left:* Representative images are shown. *Right:* Quantification of maximum number of crossings by Sholl analysis **(A)** and synaptic pairs **(B)** is shown. Data are mean of n = 13-39 cells in **(A)** and n = 14-35 cells in **(B)** from 3 independent experiments. **(C-D)** Neurons pretreated with vehicle (DMSO), 10 μM H89, 30 μM Dyngo-4a or 100 nM Torin for 30 min, then stimulated with 1 μM Iso for 30 min. Representative Western blots of 4E-BP1 phosphorylation (Thr37/46) and total 4E-BP1 are shown. Error bars represent standard error of the mean. **** = *p* < 0.0001 by two-way ANOVA with Sidak correction. ND = no drug; Veh = vehicle. Scale bars, 50 μm in **(A)** and 10 μm in **(B)**.

In parallel with the transcriptional programs, we examined the translationally regulated gene set that exhibited dependence on β2AR endocytosis. These showed striking enrichment of mRNAs related to ribosomal structure and translational processes (15/53 targets; *p* < 1.0x10^-43^, Fisher’s exact test, **Supplementary Fig. 4e**). Moreover, translation of these genes was significantly reduced by endocytic blockade (*p* = 1.0x10^-3^, Wilcoxon test). Notably, this gene set was characterized by the presence of a 5′ terminal oligopyrimidine (5′TOP) motif, a signature conferring sensitivity to mTOR-dependent translational control^32^. Prompted by this, we examined further the effects of β2AR stimulation on TOP mRNAs. Of ∼100 annotated mammalian TOP mRNAs^32^, 89 genes were detected in our Ribo-seq datasets. Although only a subset passed our threshold for classification as ‘exclusive’ translational hit, TOP mRNAs as a group showed robust induction of translation efficiency in response to isoproterenol compared to all other genes (*p* = 8.4 x 10^-22^ by Wilcoxon test, **Supplementary Fig. 5h)**^32^. These findings raised the possibility that intracellular β2AR signaling regulates neuronal mRNA translation through the mTOR pathway. Indeed, isoproterenol increased phosphorylation of the mTOR substrate 4EBP-1. This induction was blocked by inhibitors of mTOR, PKA, and dynamin (**Fig. 5c-d, Supplementary Fig. 5i-j)**. Because mTOR is classically activated downstream of the phosphoinositide 3-kinase (PI3K)/Akt and extracellular signal-regulated kinase (Erk) signaling pathways in neurons and other cell types, we next asked whether these pathways also contribute to β2AR-dependent mTOR activation. Inhibition of Akt had no detectable effect on agonist-induced 4EBP-1 phosphorylation **(Supplementary Fig. 5j)**. Inhibition of Erk modestly increased basal 4EBP-1 phosphorylation but likewise did not suppress the response to isoproterenol **(Supplementary Fig. 5k)**. These support that mTOR activation lies downstream of PKA and requires receptor endocytosis.

Lastly, because mTOR is a well-established driver of structural plasticity downstream of receptor tyrosine kinases^33,34^, we hypothesized that endosomal β2ARs similarly engage mTOR to control morphogenesis. Indeed, application of the mTOR inhibitor, Torin, blocked β2AR-induced dendritic arborization and synapse formation (**Fig. 5a-b**, blue bars). Collectively, these results indicate that intracellular β2ARs signaling coordinates gene expression through PKA/CREB and PKA/mTOR to induce dendritic and synaptic remodeling.

### Endosomal **β**2AR signaling drives neuronal remodeling in primary cortical neurons

To determine whether the signaling mechanisms identified in iNeurons extend to a primary neuronal system, we examined β2AR responses in primary cortical neurons. Similar to iNeurons, stimulation with either the synthetic β-adrenergic agonist isoproterenol or the β2AR-selective agonist salmeterol promoted dendritic arborization and synapse formation (**Fig. 6a-b)**. These structural responses were abolished by acute inhibition of receptor endocytosis with Dyngo-4a, indicating that intracellular β2AR signaling is likewise required for morphogenesis in primary neurons (**Fig. 6a-b)**.

**Figure 6.**
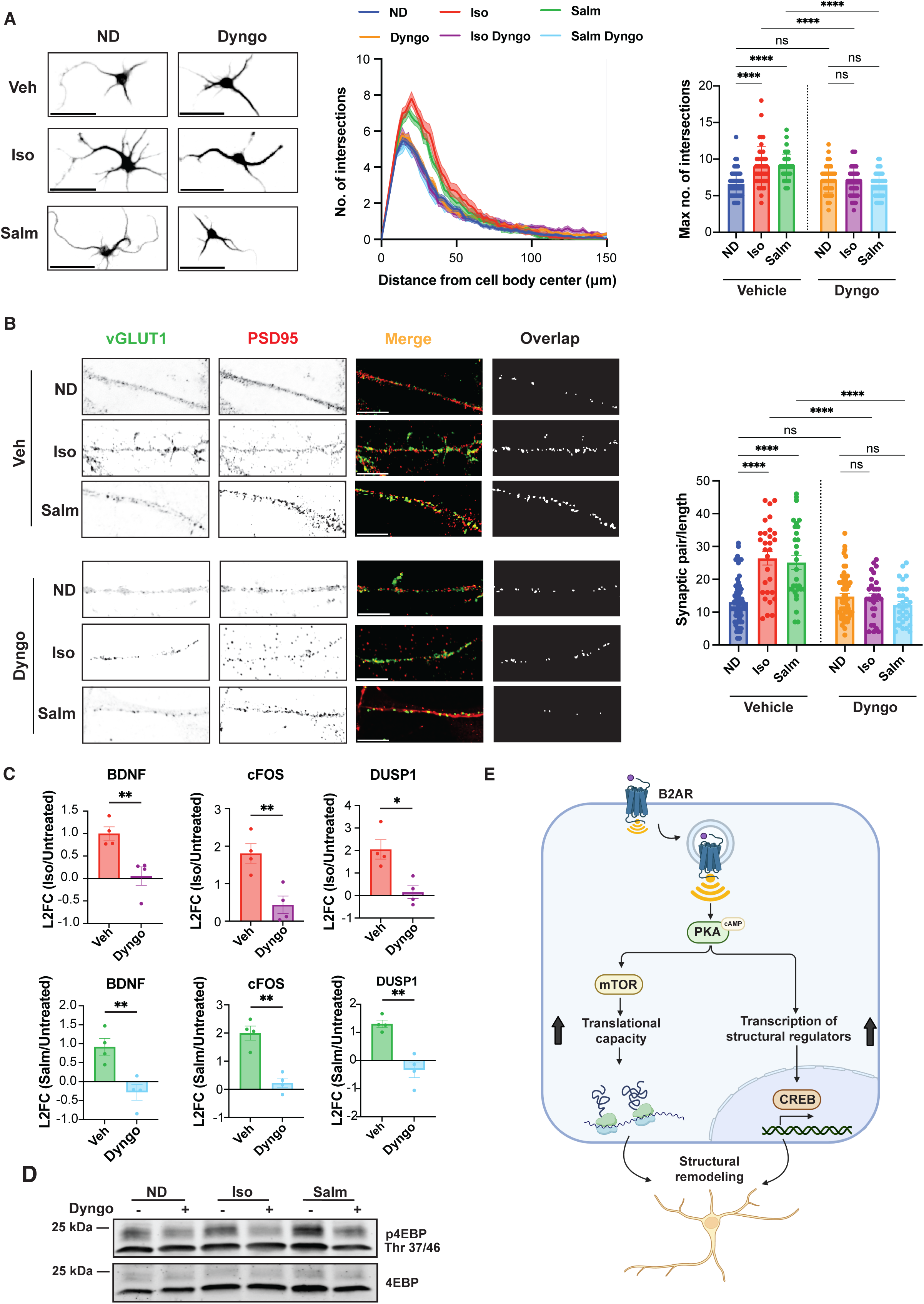
Endosomal β2AR signaling drives primary cortical neuron remodeling. **(A-B)** Analyses of dendritogenesis and synapse formation following various treatments of primary cortical neurons. Neurons were pretreated with vehicle (DMSO) or 30 μM Dyngo-4a for 30 min, then stimulated with 1 μM Isoproterenol (Iso) or 100 nM salmeterol (Salm). Neurons were fixed and stained for MAP2 **(A)** or vGLUT1/PSD95 **(B)**. *Left:* Representative images are shown. *Right:* Quantification of maximum number of crossings by Sholl analysis **(A)** and synaptic pairs **(B)** is shown. Data are mean of n = 43-45 cells for **(A)** and n = 30 cells in **(B)** from 2 independent experiments (Iso) and 3 independent experiments (Salm). **(C)** Neurons were pretreated with 30 μM Dyngo-4a for 30 min, followed by 1 μM Iso or 100 nM Salm for 2 h, RNA was extracted and *BDNF*, *DUSP1*, and *cFOS* expression was analyzed by RT-qPCR, n = 3 independent experiments. **(D)** Neurons were pretreated with 30 μM Dyngo-4a for 30 min, followed by 1 μM Iso or 100 nM Salmeterol for 1 h. Representative Western blots of 4E-BP1 phosphorylation (Thr37/46) and total 4EB-P1 are shown. **(E)** Model: Endosomal GPCR signaling elicits morphogenesis through transcriptional and translational changes that together boost the cell’s protein synthesis capacity to provide key structural effectors. Error bars represent standard error of the mean. **** = *p* < 0.0001 by two-way ANOVA with Sidak correction **(A-B)** and by unpaired Student’s *t* test **(C)**. ND = no drug; Veh = vehicle. Scale bars, 50 μm in **(A)** and 10 μm in **(B)**.

We next asked whether endosomal β2AR activity is similarly required for the induction of transcriptional and mTOR signaling. β2AR stimulation induced the expression of IEGs and the neurotrophic factor *BDNF*, and this transcriptional response was completely abolished by endocytic blockade (**Fig. 6c)**. Likewise, β2AR activation increased phosphorylation of the mTOR substrate 4EBP-1, and this response was also inhibited by Dyngo-4a (**Fig. 6d)**. Together, these results establish that the endosomal β2AR signaling pathways identified in human iPSC-derived neurons also operate in primary cortical neurons and support intracellular receptor signaling as a fundamental mechanism for coordinating neuronal gene expression and structural plasticity.

## DISCUSSION

While the classical view held that GPCRs signal primarily from the plasma membrane, a growing body of work has established that many neuronal GPCRs also signal from intracellular compartments, including endosomes, the Golgi, endoplasmic reticulum, and mitochondria^35–40^. These studies have demonstrated that intracellular receptor signaling can regulate diverse aspects of neuronal physiology, such as excitability, synaptic transmission, and metabolism. Nevertheless, how signals originating from intracellular receptor pools are decoded into distinct biochemical and cellular programs has remained incompletely understood. Our previous studies in HEK293 cells established that endosomal β2AR signaling promotes transcriptional responses and translational control^16,24,30^. Here, we show that this compartmentalized signaling logic also operates in neurons. There, it engages a substantially broader gene regulatory responses uniquely enriched for regulators of neuronal morphogenesis and promotes dendritic growth and synapse formation. These findings therefore establish not only that the spatial logic of compartmentalized β2AR signaling is conserved in physiologically relevant cell types, but also that its downstream outputs are adapted to execute cell type-specific biological programs.

Activation of endosomal β2ARs elicits two coordinated molecular responses: transcriptional induction of genes involved in neuronal development and morphogenesis together with selective enhancement of translation efficiency for mRNAs encoding components of the protein synthesis machinery (**Fig. 3, 4)**. This is consistent with a coordinated two-tiered mechanism for promoting structural plasticity driven by the internal receptor: a rapid translational response that amplifies the protein synthesis capacity of the cell coupled with a transcriptional program that supplies key structural regulators (**Fig. 6e)**. We also note that, while the primary transcriptional response was enriched for pathways associated with neuronal differentiation and morphogenesis, several additional functional categories also emerged. Among these were genes involved in feedback regulation of intracellular signaling, including enrichment of the “inactivation of MAPK activity” pathway, driven in part by induction of dual-specificity phosphatases (DUSPs) (**Fig. 3)**. Although the functional significance of these pathways was not investigated here, they likely reflect broader adaptive responses downstream of β2AR activation and represent interesting directions for future studies.

In principle, endosomal β2ARs could engage multiple downstream signaling cascades^16,22,24,41,42^. Our data identify PKA as the primary downstream effector that decodes the signal to modulate neuronal gene expression and ultimately promote neuronal morphogenesis. Specifically, we find that these processes are mediated by PKA-dependent activation of transcription through CREB and translation through mTOR (**Fig. 5, Supplementary Fig. 5)**. Both CREB and mTOR have been recognized as regulators of neuronal structural plasticity. Specifically, neurotrophic signaling through BDNF/TrkB promotes dendritic growth and protein synthesis by engaging CREB and mTOR^43,44^. Likewise, recent studies have shown that 5-HT2A receptor signaling promotes dendritic and synaptic remodeling through mTOR-dependent mechanisms^45,46^. Interestingly, we identify that the β2AR engages a mechanistically distinct route that converges on these same downstream regulators. Whereas neurotrophic and serotonergic signaling promote neuronal structural plasticity primarily through PI3K/Akt-, Erk-, and Ca²⁺-dependent pathways, endosomal β2AR signaling couples activation of both CREB and mTOR to PKA. Consistent with this, β2AR-dependent mTOR activation was insensitive to inhibition of either Akt or Erk **(Supplementary Fig. 5)**. Defining how PKA communicates with the mTOR pathway downstream of endosomal β2ARs will be a critical direction for future investigation. These observations suggest that mechanistically diverse receptor systems converge on common transcriptional and translational hubs while employing fundamentally distinct upstream signaling architectures.

An additional striking similarity among these pathways is that the receptor populations responsible for driving structural plasticity are intracellular. Specifically, axonal BDNF/TrkB signaling endosomes activate CREB and mTOR to promote dendritic growth, whereas intracellular pools of 5-HT2A receptors preferentially drive psychoplastogenic responses through mTOR-dependent signaling^44,47^. In combination with our findings, these suggest that intracellular receptor signaling represents an evolutionarily conserved organizational strategy through which neurons convert transient extracellular stimuli into durable structural plasticity.

An important remaining amechanistic question is how the GPCR signal is compartmentalized to ultimately regulate gene expression. According to our data, PKA activity remains limited across neuronal compartments when β2AR endocytosis is blocked **(Supplementary Fig. 5)**. Increasing evidence supports that compartmentalized cAMP signaling is maintained through the combined actions of phosphodiesterases (PDEs) and A-kinase anchoring proteins (AKAPs) that generate localized signaling nanodomains to restrict diffusion of second messengers^48,49^. In HEK293 cells, we have previously shown that β2AR signaling originating at the plasma membrane is buffered by higher local PDE activity relative to endosomes, thereby conferring distinct signaling properties to these receptor pools^16,30,50^. Whether analogous mechanisms establish and maintain endosomal β2AR signaling domains in neurons remains to be determined.

More broadly, our findings raise the intriguing possibility that receptor internalization itself functions as a biological gate for structural plasticity. This “spatial checkpoint” could preferentially couple stimuli of sufficient magnitude or duration to the transcriptional and translational programs that drive morphogenesis. One attractive possibility is that neurotransmitter concentration and kinetics determine when this checkpoint is engaged. Brief, low-concentration norepinephrine exposure may primarily activate plasma membrane β2ARs and favor acute physiological responses without inducing structural remodeling. In contrast, higher or more sustained adrenergic stimulation, as might occur during heightened arousal or attention, would be expected to promote receptor internalization and thereby favor long-term structural plasticity.

Altogether, the present work illuminates new spatial and biochemical mechanisms, by which GPCR signaling modulates neuronal function. Moving forward, it will be important to establish how broadly these mechanisms apply across neuronal types and circuits, and how they ultimately affect network activity and behavior.

## ACKNOWLEDGEMENTS

We thank members of the Tsvetanova lab for their thoughtful feedback on the study. All microscopy imaging was performed using resources from the Duke Light Microscopy Core Facility (LMCF) with training under Drs. Lisa Cameron and Yasheng Gao. This work was supported by the NIH (R01NS127847 to N.G.T. and 5T32GM133352 to K.L.H) and American Heart Association (25PRE1377010 to K.L.H.). Fig. 1a, Fig. 6e and Supplementary Fig.1b were created using BioRender.

## AUTHOR CONTRIBUTIONS

N.G.T. conceived and supervised the study and acquired funding. N.G.T., K.L.H., and M.J.K. designed the study and developed the experimental approaches. K.L.H. performed and analyzed the neuronal signaling, gene expression, dendritic morphogenesis, and synaptogenesis experiments, established and performed experiments in primary cortical neurons, and developed and implemented the compartment-targeted Nb80 experiments. M.J.K. optimized the RNA-seq and Ribo-seq approaches. M.J.K and K.L.H performed and analyzed the RNA-seq and Ribo-seq experiments. K.L.H. and M.E.S. performed and analyzed Western blot experiments. K.L.H. and N.G.T. prepared the figures and wrote the manuscript with input from all authors. All authors reviewed and approved the final manuscript.

## MATERIALS AND METHODS

### Chemicals and antibodies

(−)-Isoproterenol hydrochloride (Sigma-Aldrich, Cat #I6504) was purchased from, dissolved in 100 mM ascorbic acid to 10 mM stock, and used at indicated concentrations. Dyngo-4a (Abcam, Cat #120689) was resuspended in DMSO to 30 mM stock and used at 30-45 μM final concentration. 666-15 (MedChemExpress, Cat#HY-101120) was resuspended to 1 mM in DMSO and used at a final concentration of 1 μM. Torin-1 (MedChemExpress, Cat #HY-13003) was resuspended to 100 μM in DMSO and used at a final concentration of 100 nM. H89 (Cayman Chemical, Cat #10010556) was dissolved in DMSO to 10 mM and used at 10 μM final concentration. 8-bromo-cAMP (Abcam, #ab141448) was dissolved in water to 100mM and used at 100 μM final concentration. Pitstop 2 (Abcam, #ab120687) was dissolved in DMSO to 30 mM and used at 30 μM final concentration. BLU0588 (MedChemExpress, Cat #HY-153967) was dissolved in DMSO to 1 mM and used at 1 μM final concentration. Salmeterol (MedChemExpress, Cat #HY-14302) was dissolved in DMSO to 100 μM and used at 100 nM final concentration. Norepinephrine (Sigma-Aldrich, Cat #A7257) dissolved in 100 mM ascorbic acid to 10 mM stock and used at 10 μM final concentration. SCH-772944 (MedChem Express, Cat #HY-50846) was resuspended at 10 mM in DMSO and used at 10 μM final concentration. MK-2206 (MedChem Express, Cat #HY-10358) was resuspended at 1mM in DMSO and used at 1 μM final concentration.

Rabbit anti-4E-BP1 monoclonal antibody (Cell Signaling Technologies, Cat #9644 and Cat #9452L), rabbit anti-phospho-4E-BP1 monoclonal antibody (Thr37/Thr46) (Cell Signaling Technologies, Cat #2855), rabbit anti-FTH1 monoclonal antibody (Cell Signaling Technologies, Cat #4393), rabbit anti-FOS monoclonal antibody (Cell Signaling Technologies, Cat #2250T), rabbit anti-NPAS4 monoclonal antibody (Activity Signaling, Cat #AS-AB18A-100), mouse anti-beta-Actin monoclonal antibody (Santa Cruz Biotechnology Cat #sc-69879) were used at 1:1,000 for Western blots. Chicken anti-MAP2 monoclonal antibody (Phosphosolutions, Cat #1099-MAP2), rabbit anti-NGN2 polyclonal antibody (Invitrogen Cat #PA5-78556), guinea pig anti-VGLUT1 polyclonal antibody (Millipore, Cat #AB5905), rabbit anti-PSD95 polyclonal antibody (Abcam, Cat #AB18258) were used at 1:1,000 for immunofluorescent staining. Mouse anti-CREB monoclonal antibody (Cell Signaling Cat #9104) and rabbit anti-phospho-CREB monoclonal antibody (Ser133) (Cell Signaling Technologies, Cat #9198) were used at 1:100 for immunofluorescent staining. The following secondary antibodies were used at 1:10,000 for Western and immunofluorescence: donkey anti-mouse IRDye 680RD (Licor, Cat #926-68072), donkey anti-rabbit IRDye 800CW (Licor, Cat #925-32213), goat anti-guinea pig Alexa Fluor 488 (Thermo Fisher Scientific, Cat #A11073), goat anti-guinea pig Alexa Fluor 568 (Thermo Fisher Scientific, Cat #A11075), goat anti-chicken Alexa Fluor 568 (Thermo Fisher Scientific, Cat #A11041), donkey anti-rabbit Alexa Fluor 488 (Thermo Fisher Scientific, Cat #A21206), donkey anti-rabbit Alexa Fluor 647 (Thermo Fisher Scientific, Cat #A31573).

### Lentivirus production

Lentivirus was used to express ExRai-AKAR2, ExRai-AKAR2-NLS, Nb37-eGFP, Nb80-eGFP, 2xFYVE-Nb80-eGFP, and NLS-Nb80-eGFP. HEK293T were transfected with lentiviral constructs and packaging vectors (psPAX2 and VSVG) using Lipofectamine 2000 (Thermo Fisher Scientific, Cat #11668019) and OptiMEM (Thermo Fisher Scientific, Cat #31985062). Media was changed 16 h post transfection and the supernatant was harvested 72 h post transfection. The viral supernatant was concentrated using LentiX concentrator (Takara, Cat #631231), resuspended in PBS, and snap-frozen or used immediately.

### Construct cloning

The lentiviral GFP-tagged Nb37 and Nb80 constructs were described previously^30^. To generate pFUGW-ExRai-AKAR2 plasmid, AKAR2 sequence was PCR amplified from the original ExRai-AKAR2 and ExRai-AKAR2-NLS constructs^30,31^ and inserted into a digested pFUGW-GFP backbone by In-Fusion cloning (Takara Bio, Cat #638947) following recommended protocols. To generate 2xFYVE-Nb80-eGFP, 2xFYVE sequence was amplified and inserted into the pFUGW-Nb80-eGFP digested backbone by NEBuilder HiFi DNA Assembly cloning kit (NEB, Cat #E5520) following recommended protocols. To generate NLS-Nb80-eGFP, the NLS-Nb80-eGFP sequence was generated by TWIST Bioscience, PCR amplified and inserted into a digested pFUGW backbone by NEBuilder HiFi DNA Assembly cloning kit (NEB, Cat #E5520) following recommended protocols.

### iNeuron cell culture

The human iPSC line expressing Flag-tagged β2AR was previously generated and characterized in Ref ^20^. Briefly, hiPSCs expressing *Ngn2* under a doxycycline-inducible promoter integrated into the AAVS1 safe harbor locus^21^ were transduced with Flag-β2AR under ubiquitin promoter and low-expressing clones were isolated by fluorescent cell sorting. This line expresses ∼20-fold higher *ADRB2* mRNA than the parental cells, which provided sufficient dynamic range for live-cell imaging and genome-wide transcriptional and translational analyses. As previously described, comparable β2AR expression levels in HEK293 cells do not result in superphysiological signaling^20^. For PKA activity measurements, we generated stable iPSC lines expressing either ExRai-AKAR2 or NLS-ExRai-AKAR2 by transducing Flag-β2AR cells with either construct. The same stable reporter lines were used for all neuronal differentiations and experimental conditions involving these biosensors in order to minimize variability in sensor expression associated with transient transfection or viral transduction of differentiated neurons.

iPSCs were grown in mTeSR media (StemCell Technologies, Cat #100-1130) with 10 μM Rock inhibitor (MedChem Express, Cat #HY-10583-10mg) on plates with Matrigel Basement Membrane Matrix (Corning, Cat #356231) diluted in Knockout DMEM (Thermo Fisher Scientific, Cat #10829018) to 100 μg/mL. Media changes were performed daily. For differentiation into iNeurons, iPSCs were grown for 3 days in Knockout DMEM/F12 (Thermo Fisher Scientific, Cat #12660012) supplemented with 1X N2 (ThermFisher Cat #17502048), 10 ng/mL Recombinant Human NT-3 (PeproTech Cat #450-03), 2 μg/mL Doxycycline (Sigma, Cat #D3447), 10 ng/mL Recombinant Human/Murine/Rat BDNF (PeproTech, Cat #450-02), 1 μg/mL Mouse Laminin (Thermo Fisher Scientific, Cat #23017015) and 1X MEM Non-Essential Amino Acids (NEAA) (Thermo Fisher Scientific, Cat #11140050). Then, neurons were replated and grown on BioCoat PDL-coated plates (Corning, Cat #354413) in 0.5X Neurobasal-A media (Thermo Fisher Scientific, Cat #10888022) and 0.5X DMEM/F-12 media (Thermo Fisher Scientific, Cat #11330032) supplemented with 0.5X N2 supplement, 2 μg/mL Doxycycline, 10 ng/mL recombinant NT3, 10 ng/mL recombinant BDNF, 0.5X B-27 supplement (Thermo Fisher Scientific, Cat #12587010), 0.5X Glutamax (Thermo Fisher Scientific, Cat #35050-061) and 1X NEAA. On day 7, half of the culture medium was replaced with an equal amount of fresh culture medium without doxycycline. On day 14, half of the medium was again removed, and fresh culture medium without doxycycline was added at twice that volume.

### Primary neuron dissections and culture

Rats were maintained in Duke’s Division of Laboratory Animal Research (DLAR) Vivarium facility. All experiments with animals have complied with ethical regulations for animal testing and research in accordance with federal regulations mandated by the Institutional Animal Care and Use Committee (IACUC). DUIACUC Protocol #: A006-24-01.

Embryonic cortical neurons were obtained from E19 CD-Sprague Dawley Rats from Charles River Laboratory. The pregnant rat was euthanized, and embryonic brains were removed. The cerebral cortex was dissected out from both hemispheres and carefully meninges removed from cortex. For the dissociation, tissue in cold HBSS was spun down, aspirated and warmed dissociation solution was added for 10 min at 37°C (100 U Papain (Worthington Biochemical, Cat #LS003126), 3.2 mg L-cysteine (Millipore Sigma, Cat #C7880) in 5 mL of HBSS). Tissue was spun again, aspirated and resuspended in warmed trypsin neutralizing solution (Lonza, Cat #CC-5002) and this step was repeated. Tissue was then resuspended in cortical plating media (Basal Media Eagle (Sigma-Aldrich, Cat #B1522), 10% calf serum (Millipore Sigma, Cat #12138C), 1% Pen/strep (Gibco, Cat #15140122), 1% Glutamax (ThermoFisher Scientific, Cat #35050061) and 6.67ml 45% glucose) and triturated 10-15x to break up the pellet. Cells were spun down and media was aspirated and resuspended in fresh cortical plating media before being counted by hemocytometer. Neurons were plated at 20k/well for IF experiments on 12 mm PDL-laminin-coated acid-etched coverslips (described below), 500k/well on PDL-coated 12-well plates or 1M/well in PDL-coated 6-well plates for the signaling assays. After 4 hours, cortical plating media was aspirated and replaced with NB/B27 media (Neurobasal (Gibco, Cat #21103-049), 2% B27 supplement (Thermo Fisher Scientific, Cat #17504044), 1% pen/strep, 1% Glutamax). For cultures longer than DIV7, 1% FBS was added. Cells were half-fed every 4 days until collection. For RT-qPCR and western blot experiments of primary cultures, media containing 1% FBS, was changed to media with no FBS on DIV20 and treated on DIV21.

### RNA-seq and Ribo-seq sample prep

iNeurons were plated and differentiated at ∼1.2 million cells per well in BioCoat PDL-coated 6-well plates (Corning Cat, #354413) and 6 wells were pooled per biological replicate per condition. At 21 days of differentiation, cells were treated with either 45 μM of Dyngo-4a or vehicle (DMSO) for 20 min to ensure uniform and robust inhibition of dynamin, and then stimulated with 1uM Isoproterenol for 2 h. Cells were then washed with ice-cold PBS and lysed in freshly prepared RNase-free Lysis buffer (20 mM Tris-HCl pH 7.4, 150 mM NaCl, 5 mM MgCl_2_, 1 mM DTT, 1% Triton X-100, 100 µg/mL cycloheximide, 25 U/mL TURBO DNase (Thermo Fisher Scientific, Cat #AM2238). Cells were triturated 10 times by passing through a 25G needle and clarified by centrifuging at 20,000x*g* for 10 minutes at 4°C. Each lysate was split for subsequent RNA-seq and Ribo-seq library preparation. For RNA-seq, total RNA was isolated using Zymo RNA Clean & Concentrator-25 kit (Genesee, Cat #11-353) and 1 µg total RNA was used as input for library generation using the NEBNext Poly(A) mRNA Magnetic Isolation Module (New England Biolabs, Cat #E7490) and NEBNext Ultra II RNA Library Prep Kit for Illumina (New England Biolabs, Cat #E7775). For Ribo-seq libraries, an aliquot of lysate containing around 10 µg was used as input. Ribo-seq samples were generated from these lysates following previously published protocols^24^.

### RNA-seq and Ribo-seq analysis

Raw data files were trimmed by removing adaptors, sequences aligning to ncRNA, tRNA, and rRNA were removed using Bowtie (version 2.3.5), and the remaining reads were aligned to the Genome Reference Consortium Human Build 38 (GRCh38) using STARaligner (version 2.7.5). RPF reads were extracted using length reads of 26-34 nucleotides using STARaligner FeatureCounts (Subread version 1.6.3).

R was used for building an annotation hub database and DESeq2 (version 3.16) with Wald test was implemented for differential expression analysis of raw reads^51^. Transcriptional targets were defined as RNAs with significant changes from the RNA-seq dataset using an adjusted *p*-value cutoff of 0.01 and an absolute gene log₂-fold change (Isoproterenol-treated/untreated) > 0.5. ΔTE values were computed by incorporating unstimulated RNA samples as an additional covariate in the DESeq input, as detailed in Ref^52^, and z-scores were calculated based on ΔTE distribution for each gene. Significant RPF changes for the Riboseq analysis were determined using adjusted *p*-value of 0.05 as cut-off. Genes were categorized as “Forwarded” if they showed significant RNAseq and RPF changes. “Exclusive” genes were defined as those with significant RPF changes, ΔTE values at least 1.5 SD from the mean in the same direction, and no significant RNA change (RNA-seq *p*-adj > 0.01). “Buffered” genes were classified as those with significant RNA changes, non-significant RPF changes, and ΔTE ≥ 1.5 SD below the mean in the same direction as the RNA change.

Transcriptional and translational hits were further categorized as driven by endosomal or plasma membrane signaling based on their dependence on endocytosis. For this, multiple unpaired *t-*test analyses were carried out comparing the RNAseq log₂-fold change values (Transcriptional targets) or ΔTE values (Exclusive translational targets) between vehicle- and Dyngo-4a-treated conditions. Cut-offs were established based on FDR using the two-stage step-up method of Benjamini, Krieger and Yekutieli. Targets with significant Dyngo-4a effects (FDR < 0.1) were designated as ‘endosomal’ hits; the remaining genes were classified as ‘plasma membrane’ hits.

For quality control metrics, Ribotish^53^, samtools^54^ and Ribotoolkit^55^ R packages were used for feature count analysis and periodicity. The package bigPint^56^ was used to calculate Pearson correlation coefficients between drug-treated conditions and library prep replicates. GOrilla GO^57^ was used for gene ontology analysis and SRplot^58^ was used to generate GO dot plot graphs.

### Spinning disk confocal imaging and analysis

The Andor Dragonfly Spinning Disk Confocal microscope equipped with 405 nm, 488 nm, 561 nm, 637 nm diode lasers, CO_2_- and temperature-controlled chamber was used for live- and fixed-cell imaging experiments and Fusion software was used for data collection.

For fixed cell microscopy-based analyses, neurons were differentiated for 21 days on PDL- and laminin-coated acid-etched glass coverslips. Following neuronal differentiation, cells were washed with PBS and then fixed at room temperature using 4% paraformaldehyde diluted in a 4% sucrose/PBS solution for 20 min. Coverslips were then washed twice in PBS, followed by 15 min in 50 mM NH_4_Cl solution and a 5-min incubation in 0.1% Triton-X 100/PBS solution. Primary antibodies were diluted in a 1% BSA/0.1% Triton-X/PBS solution and incubated with the coverslips for 2 h at room temperature or overnight at 4°C. Coverslips were washed 3 times in a 1% BSA/0.1% Triton-X/PBS solution and incubated for 30 min with secondary antibody. Coverslips were mounted with Prolong Gold antifade reagent with DAPI (Invitrogen, Cat #P36935) and stored at 4°C.

For dendritogenesis experiments, neurons were plated at ∼20k cells/well on coverslips placed in 24-well plates and grown for 3 days. Immature neurons were used for this analysis for ease of tracing and fully capturing dendritic complexity before axons get too long due to the rapid differentiation of iNeurons consistent with common practices^44,47^. On DIV3, half the media was removed and kept as “pre-conditioned media”. In the remaining media, cells were treated with 30 μM Dyngo-4a, 10 μM H89, 1 μM BLU0588, 100 nM Torin, or 1 μM 666-15 for 30 min, then stimulated with 1 μM Isoproterenol, 100 μM 8-Bromo-cAMP, 100 nM Salmeterol, or 10 μM Norepinephrine for 1 h. Following agonist treatment, the media was aspirated and replaced with the saved preconditioned drug-free media. Neurons were then fixed using the protocol above at DIV6 staining with anti-MAP2 primary antibody for dendrite visualization. Coverslips were imaged at 20x to allow for full visualization of neuron lengths. For best image analysis, regions of the slides where neurons were seeded at low confluency were captured. For dendrite quantification, the paint tool in ImageJ (Version 1.54p) was used to isolate cells from background fluorescence. Color thresholding was used to create a mask for the MAP2 fluorescence channel which highlights the fluorescence making it easier to be converted to binary for Sholl Analysis with the SNT plugin in ImageJ^59,60^. The center of the neuron was marked as the middle of the cell body and the shell radius was kept the same for all neurons. The intersection output from the Sholl profile was captured from the cell body to the furthest dendrite tip and plotted across conditions and the max number of dendrites for each cell was then plotted.

For synapse pair formation experiments, neurons were plated at ∼30k cell/well on laminin- and PDL-coated glass coverslips placed in 24-well plates and differentiated for 21 days. On DIV21, neurons were pretreated with inhibitor for 30 min, then stimulated with 1 μM Isoproterenol for 24 h. Following that, cells were fixed using anti-vGLUT1 and anti-PSD95 antibodies for pre- and post-synaptic protein visualization, respectively. MAP2 staining was also used for whole-neuron visualization. Slides were then imaged at 100x and analyzed with ImageJ. The fluorescent channels for vGLUT1 and PSD95 were deconvoluted for puncta visualization and merged, and color thresholding of the merged channel fluorescence was isolated and converted into a mask. The anti-MAP2 channel was used for ensuring 35 micron length was used for each neuron to normalize across conditions with the Line Tool in ImageJ. Number of puncta from the mask was recorded and plotted.

For nanobody imaging, neurons were differentiated at ∼500k on matrigel-coated glass-bottom imaging dishes in Neurobasal-A without phenol red (Thermo Fisher Scientific, Cat #12349015) and DMEM/F12 without phenol Red (Thermo Fisher Scientific, Cat #21041025). Neurons were transduced with lentiviral constructs (eGFP-Nb37 or eGFP-Nb80) for three days. Alexa-647 conjugated M1 mouse anti-flag antibody was added directly to the medium for 5 min to label cell-surface Flag-tagged β2AR, and neurons were stimulated as described and imaged before stimulation and after 15 min of ligand. This same protocol was also used for live-cell imaging of uninfected neurons pre-incubated with 30 μM Dyngo-4a for 30 min following 1 μM Isoproterenol stimulation for 15 min. Line scan analysis was used in ImageJ and plotted for both channels. For each channel, background was subtracted and an identical ROI was drawn around the cell. Coloc 2 in ImageJ was then used to calculate the Mander’s coefficients, which were plotted to represent the degree of overlap between the receptor and nanobody signals.

For imaging of the ExRai-AKAR2 sensors, iPSCs were infected with lentiviral constructs and differentiated on matrigel-coated glass bottom imaging dishes for live-cell analysis. Pitstop 2 was used instead of Dyngo-4a to inhibit endocytosis, because Dyngo-4a exhibits fluorescence in the green channel, which interferes with imaging-based readouts. Neurons were pre-treated for 20 min with 30 μM Pitstop or vehicle (DMSO). Next, 3 baseline reads were taken every 30 sec, neurons were stimulated with 10 nM Isoproterenol to avoid sensor saturation^30^ and imaged for every 30 sec for 10 min total. Fluorescence was quantified by defining ROIs for individual cells, measuring signals from the 480 nm channel in each frame, and determining ΔF/F values by the average of the baseline values. Based on these, area under the curve values were calculated.

For CREB and phospho-CREB IF analyses, neurons were plated at ∼30k/well on laminin- and PDL-coated glass coverslips placed in 24-well plates and differentiated for 21 days. On DIV21, cells were pre-treated with 30 μM Pitstop 2 or 10 μM H89 or vehicle (DMSO) for 30 min, followed by 1 μM Isoproterenol or no drug for 30 min. Cells were then fixed as described above and stained with 1:100 of pCREB and CREB overnight. The next day, coverslips were incubated with secondary antibody for 30 min. Fluorescence was quantified in ImageJ by defining ROIs for individual cells, measuring signals from both channels and then taking the ratio of the pCREB signal over the CREB signal for each cell. For live-cell endocytosis analysis involving EEA1, neurons were plated on Matrigel-coated imaging dishes and 3 days, then transfected with 700 ng of EEA1-dsRED using Lipofectamine 3000 (Invitrogen, Cat #L3000008) following recommended protocols. After 24 hours, neurons were imaged following labeling of cell-surface receptors as described and imaged after 15 min stimulation with 1 μM Isoproterenol.

### ELISA cAMP

Neurons were plated in 12-well BioCoat PDL-coated plates (Corning, Cat #354413) for 21 days at ∼250k cells/well, then pretreated with vehicle (DMSO) or 30 µM Dyngo-4a for 30 min. Neurons were stimulated for 1 µM Isoproterenol for 5 min or 15 min and the Cayman ELISA cAMP assay (VWR, Cat #75817–364) was used to quantify cAMP, following manufacturer protocols. All values were normalized to the total protein concentration of the respective sample.

### RT-qPCR

Neurons were differentiated for 21 days in 12-well BioCoat PDL-coated plates (Corning, Cat #354413) at ∼250k cells/well (iNeurons) or 500k cell/well (primary cortical neurons), pretreated with inhibitor or vehicle (DMSO) for 30 min, then stimulated with agonist for 2 h. RNA was extracted using QIAshredder (Qiagen, Cat #79656) and the Qiagen RNAeasy RNA-isolation kit (Qiagen Cat #74106) and reverse transcription was performed using SuperScript II Reverse Transcriptase (Thermo Fisher Scientific, Cat #18064014) or LunaScript Reverse Transcriptase (NEB, Cat #E3010) following manufacturer protocols. For quantitative PCR, Power SYBR Green PCR MasterMix (AppliedBiosystems, Cat #4364344) was used with the following human primers: *GAPDH Forward: GTCTCCTCTGACTTCAACAGCG; GAPDH Reverse: ACCACCCTGTTGCTGTAGCCAA; FOS Forward: GCCTCTCTTACTACCACTCACC; FOS Reverse: AGATGGCAGTGACCGTGGGAAT; FTH1 Forward: TGAAGCTGCAGAACCAACGAGG; FTH1 Reverse: GCACACTCCATTGCATTCAGCC; NPAS4 Forward: TGGCTCTACTGGACATCTCCGA; NPAS4 Reverse: CGGTAGTGTTGAGCAGAAGCGT; BDNF Forward: CATCCGAGGACAAGGTGGCTTG; BDNF Reverse: GCCGAACTTTCTGGTCCTCATC; UNCX Forward: AGAAGGCGTTCAACGAGAGCCA; UNCX Reverse: CGTGTTCTCCTTCTTCTTCCAC; CGA Forward: TCCATTCCGCTCCTGATGTGA; CGA Reverse: CGTCTTGGACCTTAGTGGAG; DUSP1 Forward: CAACCACAAGGCAGACATCAGC; DUSP1 Reverse: GTAAGCAAGGCAGATGGTGGCT*. For qRT-PCR in primary neurons the following rat primers were used: GAPDH Forward: *CAACTCCCTCAAGATTGTCAGCAA*; *GAPDH Reverse GGCATGGACTGTGGTCATGA*; *BDNF Forward*: *CGAAGAGCTGCTGCTGGATGAG*; BDNF Reverse *ATGGGATTACACTTGGTCTCG*; *DUSP1 Forward CTCCCGAGTTCCACTGAGTT*; *DUSP1 Reverse: AGAGTCCTTTCCCTTCTGCC; cFOS Forward*: *AGGCTGACTCCTTCCCTAGC; cFOS Reverse: ATGCACCAGCTCAGTCAGTG* All gene levels were normalized to the housekeeping gene *GAPDH*. The CFX-384 Touch Real-Time PCR System equipped with CFXMaestro software (BioRad) was used for analysis.

### Western Blot Analysis

Neurons were plated on BioCoat PDL-coated 6-well plates (Corning, Cat #354413) at 750k/well for iNeurons or 1M/well for primary neurons and treated on DIV20-22 as specified, washed with PBS and then lysed with ice-cold RIPA buffer (Sigma, Cat #R0278) containing phosphatase inhibitors (Sigma, Cat #P0044), protease-inhibitor cocktail (Sigma, Cat #P8340), and 1 mM PMSF (Sigma, Cat #329-98-6) on a shaker for 15 min at 4°C before. Cells were scraped off the wells and transferred into pre-chilled microcentrifuge tubes. After cold centrifugation, the supernatant was collected and protein concentration was measured using the Peirce Protein Assay kit (Thermo Fisher Scientific, Cat #23227). Equal amounts of total protein were then boiled in 4X Laemmli buffer with 10% 2-mercaptoethanol for 5 min at 90°C. Protein was loaded into Mini-PROTEAN TGX Stain-Free 4–15% gels (BioRad, Cat #4561094). Gels were transferred to PVDF membrane (Sigma, Cat #IPFL00010). Membranes were blocked with Intercept Blocking buffer/5% BSA (Licor, Cat #927-60001) for 1 h. Primary antibodies were added to the Intercept Blocking buffer and incubated overnight with shaking at 4°C. After washing in Tris Buffer Saline (TBS) solution with 1% Tween (Sigma, Cat #P9416-50ML), secondary antibodies were diluted in the Blocking buffer and incubated for 1 h at RT with shaking. The membrane was washed with TBS and imaged using the Odyssey imager system equipped with ImageStudio software (LICOR).

