## Supplementary Figures for "GPCR endocytosis rewires neuronal gene expression and cellular architecture"

**This PDF file includes:**

Figures S1 to S5

**Other supplementary information for this manuscript includes:**

Datasets S1 to S3

### SUPPLEMENTARY FIGURES

#### Supplemental Figure 1

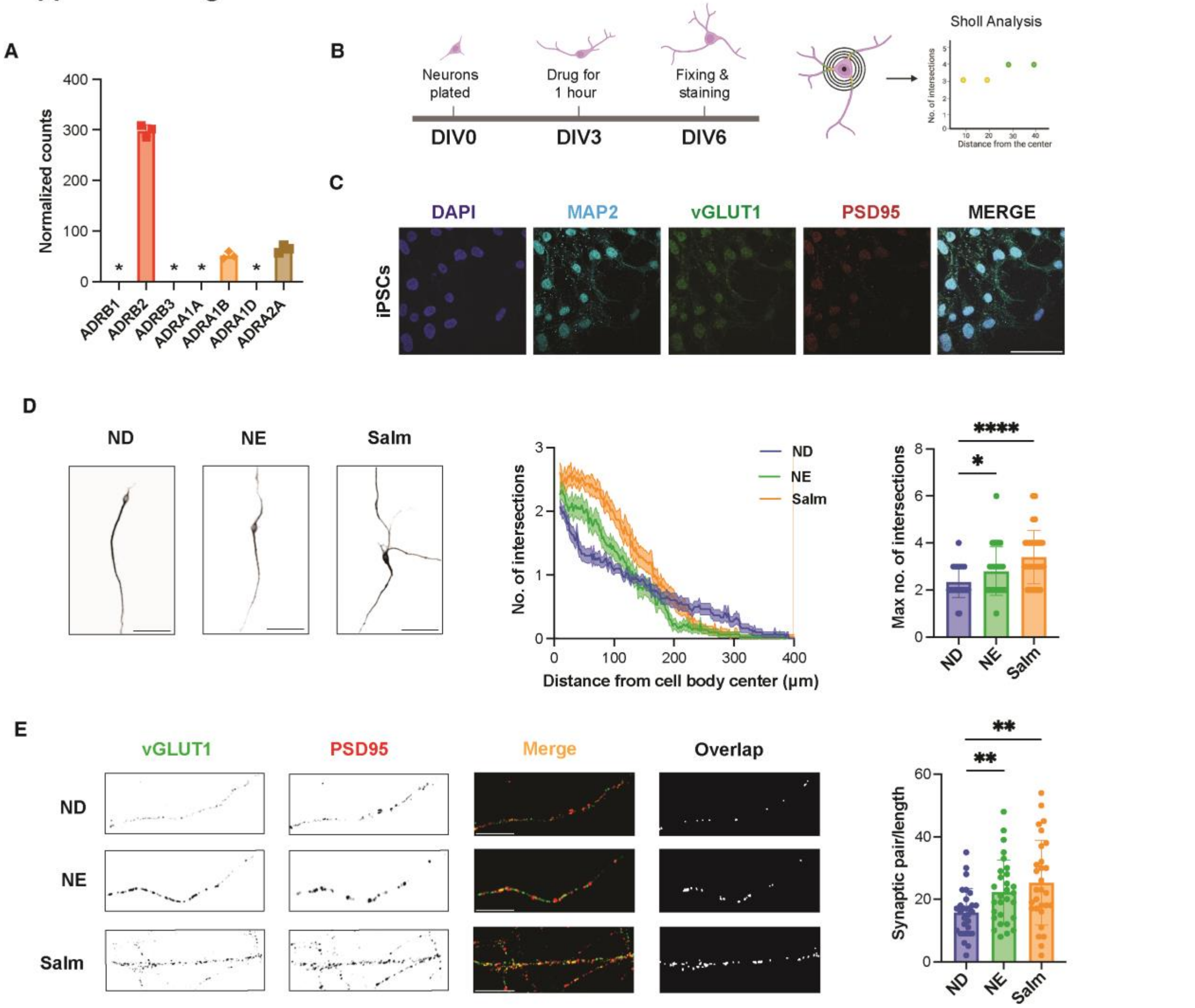


**Supplementary Figure 1. Activation of β2AR with distinct agonists induces neuronal morphogenesis. (A)** Normalized counts showing mRNA abundance of adrenergic receptor genes from RNA-sequencing in iNeurons. * = no counts detected, n = 3 independent experiments. **(B)** Schematic of dendritogenesis using Sholl analysis. iNeurons treated on post-differentiation Day 3 were treated for 1 h, then the media was replaced, and neurons were fixed and stained on Day 6. Sholl analysis marks the center of the cell body and outlines concentric rings until the final dendrite is detected to quantify number of intersections between the cell and the concentric circle. **(C)** Undifferentiated iPSCs stained with MAP2, vGLUT1, PSD95 antibodies. **(D-E)** *Left:* Representative images of dendritogenesis and synapse formation analyses are shown. iNeurons were fixed and stained for MAP2. **(D)** or vGLUT1/PSD95 **(E)**. iNeurons were stimulated with 100 nM Salmeterol (Salm) or 10 μM Norepinephrine (NE). *Right:* Quantification of maximum number of crossings by Sholl analysis **(D)** and synaptic pairs **(E)**. Data are mean of n = 30 cells in **(D)**

and n = 30 cells in **(E)** from 3 independent experiments. Error bars represent standard error of the mean. . **** = $p < 0.0001$, *** = $p < 0.001$, ** = $p < 0.01$, * = $p < 0.05$ by one-way ANOVA with Tukey correction. ND = no drug. Scale bars, 50 μm in **(D)** and 10 μm in **(C,E)**.

Supplemental Figure 2

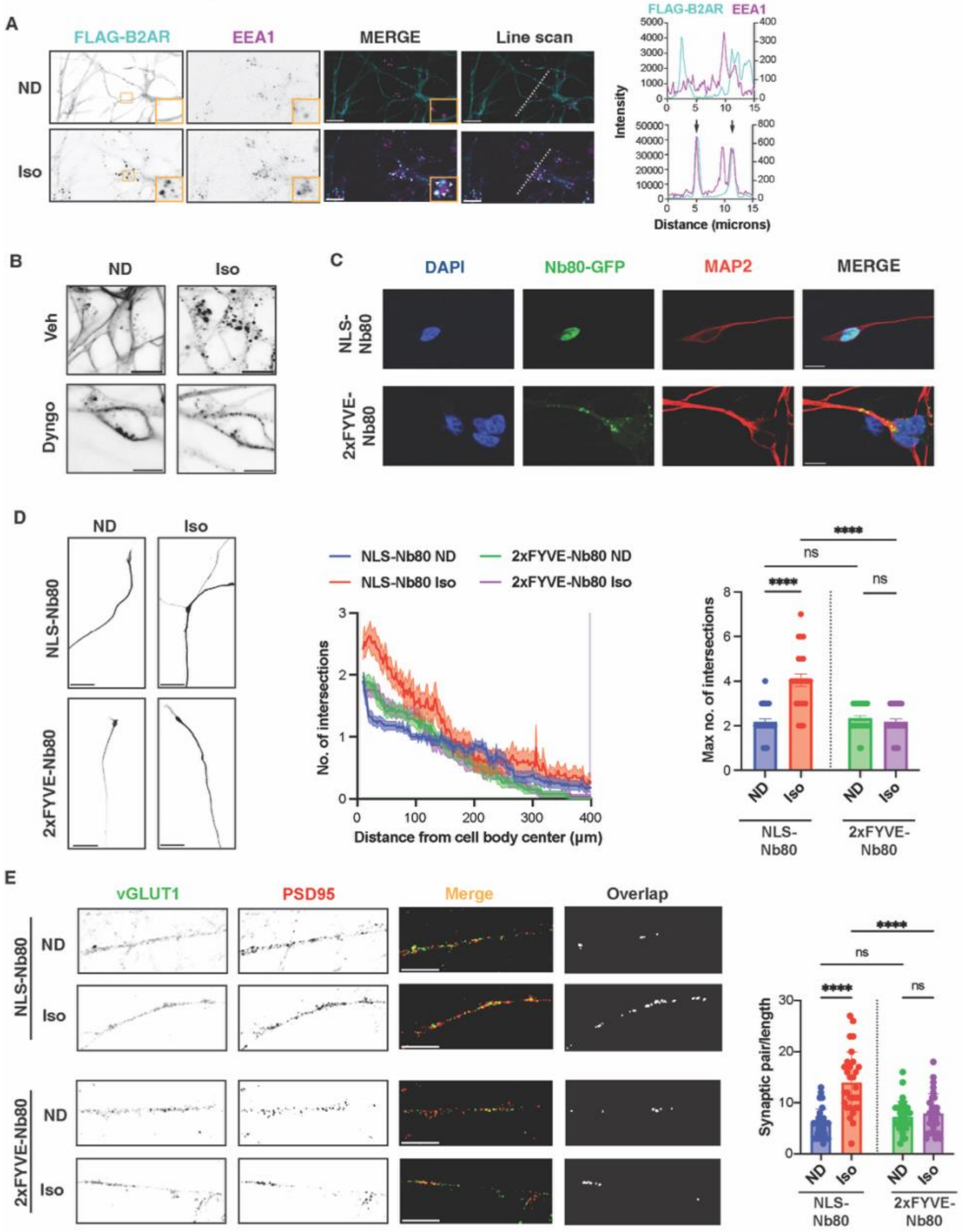


**Supplementary Figure 2. Intracellular β2AR activation drives neuronal structural responses. (A)** *Left:* Representative images of iNeurons treated with 1 µM Isoproterenol (Iso) for 15 min showing co-localization of early endosome marker EEA1 (magenta) and flag-β2AR (cyan). These are representative from n = 10 cells analyzed from 2 independent experiments. *Right:* Line scan analyses of colocalization between the β2AR and EEA1. White dotted line through the geometric center of the cell was used to analyze fluorescence intensity of each channel separately. Arrows indicate colocalization**.** Inset shows zoomed in image of region of colocalization. **(B)** Live-cell imaging of flag-β2AR localization in iNeurons pretreated with vehicle (DMSO) or 30 µM Dyngo-4a for 30 min before and after stimulation with 1 µM Iso for 15 min. Representative images were selected from n = 12-15 cells from 3 independent experiments. **(C)** Representative images of localization of NLS-Nb80-GFP and 2xFYVE-Nb80-GFP in iNeurons, co-stained with DAPI and MAP2. Data shown are representative from n = 30-40 cells from 3 independent experiments. **(D-E)** *Left:* Representative images of dendritogenesis and synapse formation analyses are shown. Neurons were fixed and stained for MAP2 **(D)** or vGLUT1/PSD95 **(E)**. iNeurons were stimulated with 1 μM Iso. *Right:* Quantification of maximum number of crossings by Sholl analysis **(D)** and synaptic pairs **(E)**. Data are mean of n = 30 cells in **(D)** and n = 30 cells in **(E)** from 3 independent experiments. Error bars represent standard error of the mean. **** = $p < 0.0001$ by two-way ANOVA with Sidak correction. ND = no drug. Scale bars, 50 μm in **(D)** and 10 μm in **(A, B, C, E)**.

Supplemental Figure 3

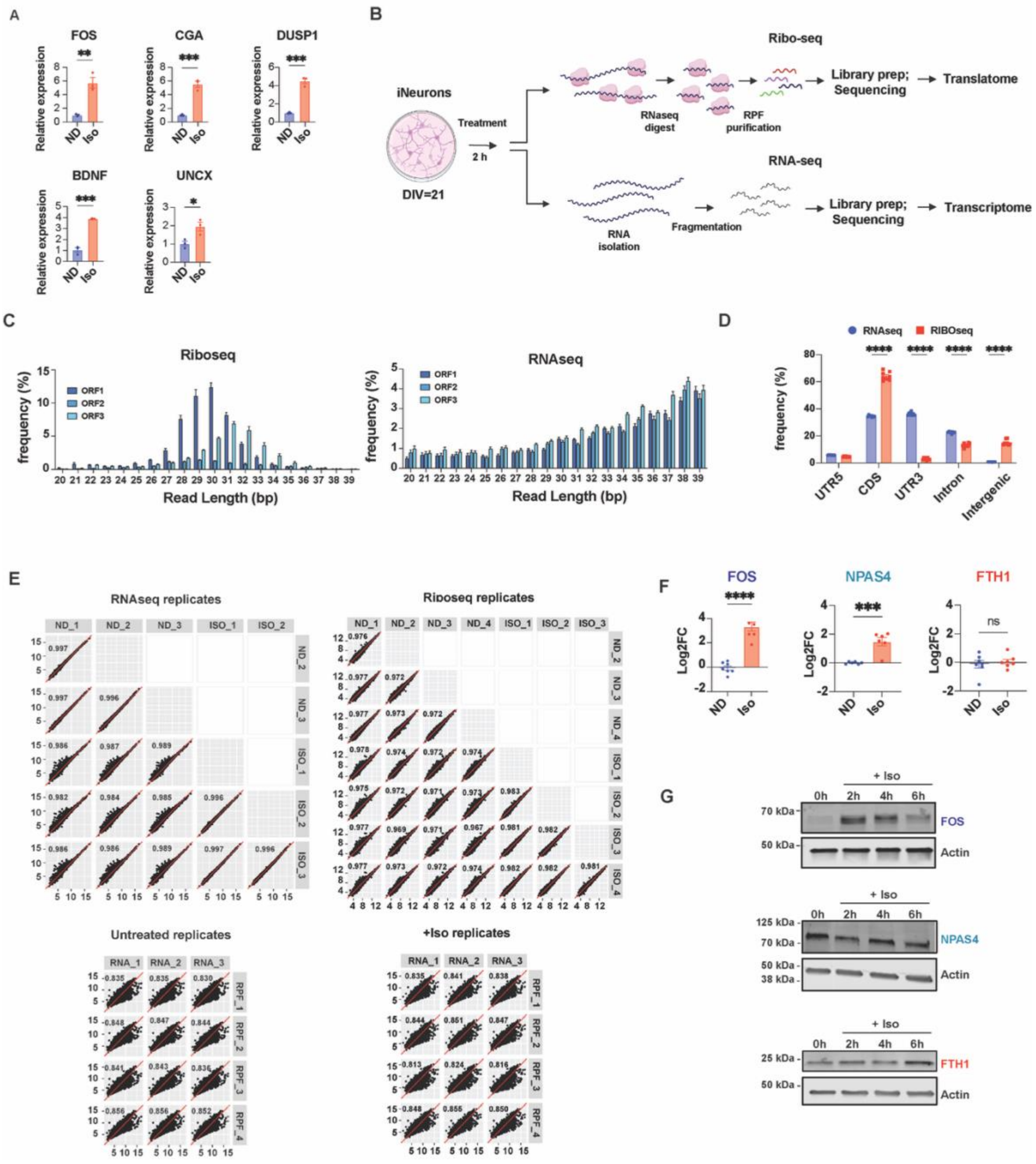


**Supplementary Figure 3. Neuronal transcriptional and translational responses to β2AR activation. (A)** RT-qPCR analysis of mRNA abundance levels for β2AR transcriptional targets encoding IEGs (*FOS, DUSP1, CGA*) and factors with known roles in dendritogenesis, axon guidance and synapse formation (*BDNF, UNCX*). iNeurons were treated with vehicle (no drug, 'ND') or 1 µM Isoproterenol (Iso) for 2 h. mRNA levels for each target were normalized to that of the housekeeping gene *GAPDH*. Data are mean of n = 3 independent experiments. **(B)** Schematic of the RNA-seq/Ribo-seq workflow to analyze the global neuronal transcriptional and translational responses to β2AR activation. **(C)** Read length distribution and periodicity of Ribo-seq (*top*) and RNA-seq (*bottom*) samples. Read length distributions were calculated using Ribotoolkit[55] and samtools[54] respectively. Periodicity was evaluated using Ribo-TISH[53]. **(D)** Genomic feature type of Ribo-seq (*top*) and RNA-seq reads (*bottom*) mapped with Ribotoolkit. **(E)** Scatter plots of correlation between Ribo-seq and RNA-seq replicates. iNeurons were treated with vehicle (ND) or 1 μM Isoproterenol (Iso) for 2 h. **(F-G)** Validation of mRNA and protein levels for select β2AR targets identified by Ribo/RNA-seq. FOS- forwarded target, NPAS4- buffered target, FTH1- exclusive translational target **(Dataset S2)**. **(F)** mRNA abundance was measured by RT-qPCR following stimulation with 1 μM Iso for 2 h and levels of each gene were normalized to *GAPDH*. Data are $\log_2$-fold change (ND/Iso) mean of n = 7 independent experiments. **(G)** Representative Western blots showing protein levels relative to beta actin (loading control). iNeurons were stimulated with 1 μM Iso for indicated times. Error bars represent standard error of the mean. **** = $p < 0.0001$, *** = $p < 0.001$, ** = $p < 0.01$, * = $p < 0.05$ by unpaired Student's *t*-test in **(A)** and **(F)** or two-way ANOVA with Sidak correction in **(D)**.

### Supplemental Figure 4

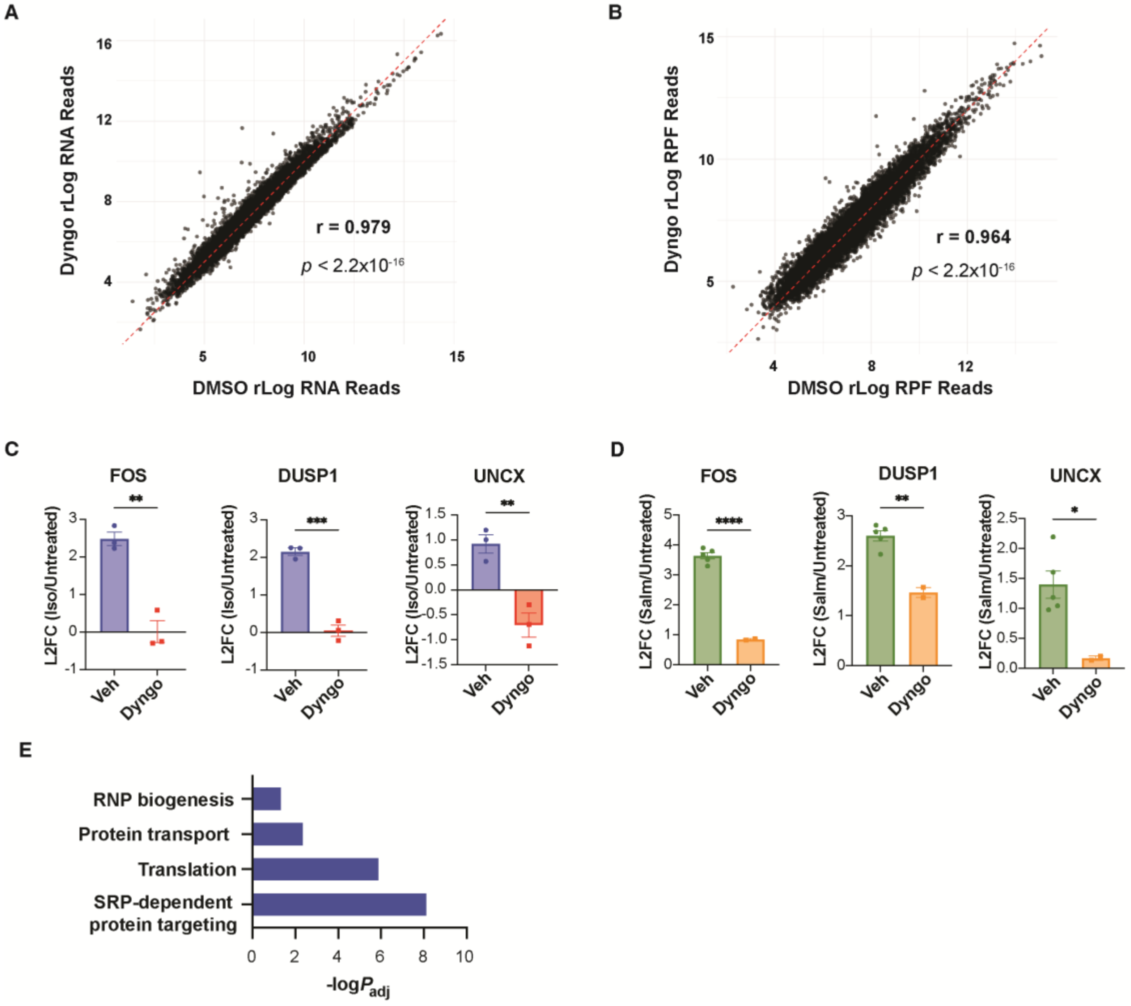


**Supplementary Figure 4. Analysis of the role of endosomal β2AR signaling in gene expression regulation. (A-B)** Effects of treatment with Dyngo-4a on basal mRNA in **(A)** and RPF read abundance in **(B)**. Scatter plots depict Pearson correlation between samples treated with vehicle (DMSO) and 45 μM Dyngo-4a. **(C-D)** RT-qPCR validation of the effects of endocytic blockade on β2AR-dependent transcriptional induction of select IEGs and morphogenesis factors. iNeurons were pretreated with vehicle (DMSO, 'Veh') or 30 μM Dyngo-4a for 30 min, then stimulated with 1 μM Isoproterenol (Iso) **(C)** or 100 nM Salmeterol (Salm) **(D)** for 2 h. mRNA levels for each target were normalized to that of the housekeeping gene *GAPDH*. Data display mean $\log_2$-fold change (agonist/no drug) values from n = 3 independent experiments for Iso and n = 2-5 independent experiments for Salm. **(E)** Gene ontology categories enriched among the exclusive translational targets that showed dependence on endocytosis **(Fig. 4** in green, **Dataset S2)**. No significant GO categories were identified for translational targets classified as 'PM'-dependent **(Fig. 4** in yellow, **Dataset S2)**. Error bars represent standard error of the mean. **** = $p < 0.0001$, *** = $p < 0.001$, ** = $p < 0.01$, * = $p < 0.05$ by unpaired Student's *t*-test in **(C-D)**.

### Supplemental Figure 5

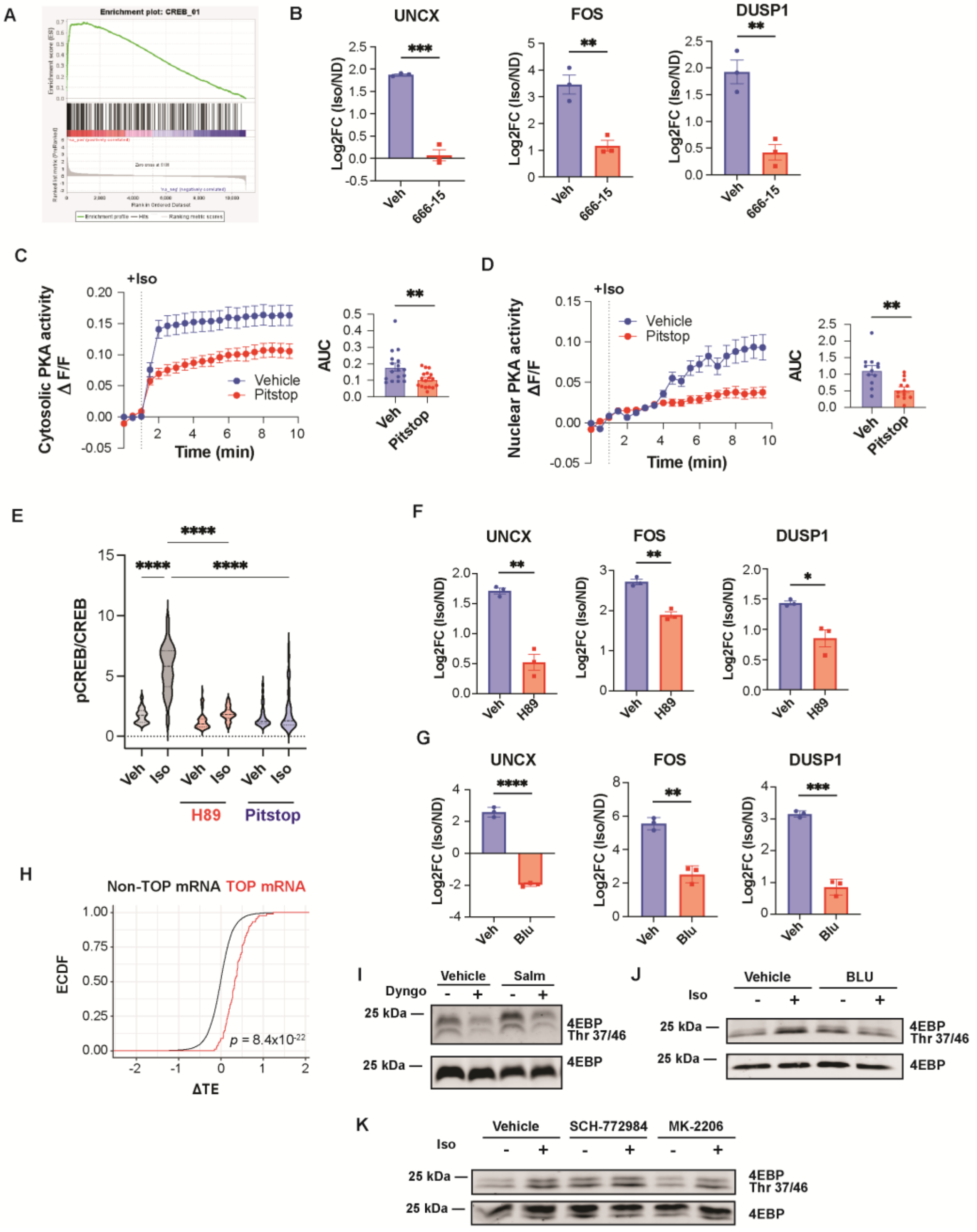


**Supplementary Figure 5. Characterization of the molecular pathways driving endosomal β2AR-dependent gene regulation. (A)** Gene Set Enrichment Analysis (GSEA)[27,28] shows enrichment of CREB targets in RNA-seq dataset ranked by averaged $log_2$-fold change (Isoproterenol/no drug) values. **(B)** iNeurons were pretreated with vehicle (DMSO) or 1 μM 666-15 for 30 min, then stimulated with 1 μM Isoproterenol (Iso) for 2 h, transcript levels were measured by RT-qPCR and normalized to *GAPDH*. Data display mean $log_2$-fold change (Iso/no drug) values from n = 3 independent experiments. **(C-D)** Real-time detection of cytosolic **(C)** and nuclear PKA activity **(D)** in neurons expressing Exrai-AKAR2 or Exrai-AKAR2-NLS sensors, respectively. Cells were pretreated with vehicle (Veh) or 30 μM Pitstop for 30 min before stimulation with 10 nM Iso (indicated by the dashed line). The Δ*F*/*F* ratio was calculated from images taken every 30 s for 10 min and the total cytosolic or nuclear PKA activity was quantified by calculating the area under the curve (AUC). Data are mean of n = 12 cells (NLS) or n = 17 cells (Cyto) from 3 independent experiments. **(E)** iNeurons pretreated with vehicle, 30 μM Pitstop or 10 μM H89 for 30 min were then stimulated with 1 μM Iso for 30 min. Phosphorylation of CREB (Ser133) was evaluated by immunofluorescence staining and is displayed as fraction of total CREB. Data are mean of n = 29-66 cells from 3 independent experiments. **(F-G) i**Neurons were pretreated with vehicle, 10 μM H89 **(F)** or 1 μM BLU0588 (BLU) **(G)** for 30 min, then stimulated with 1 μM Iso for 2 h and analyzed by RT-qPCR. Data are mean $\log_2$-fold change (Iso/no drug) values from n = 3 independent experiments. **(H)** ECDF plot of gene ΔTE value distribution in response to stimulation with 1 μM Iso for 2 h. In black is shown the distribution of mRNAs without annotated TOP motifs, in red- the distribution of genes containing TOP motifs[35]. **(I)** iNeurons were pretreated with vehicle (DMSO) or 30 μM Dyngo-4a for 30 min, followed by 100 nM Salmeterol (Salm) for 30 min. Representative images from western blots of 4E-BP1 phosphorylation (Thr37/46) are shown from 3 independent experiments. **(J)** iNeurons were pretreated with vehicle (DMSO) or 1 μM BLU0588 for 30 min, followed by 1 μM Iso for 30 min. Representative images from western blots of 4E-BP1 phosphorylation (Thr37/46) are shown. **(K)** iNeurons were pretreated with vehicle (DMSO), 10 μM SCH-772984 for 1 h, or 1 μM MK-2206 for 30 min, followed by stimulation with 1 μM Iso for 30 min. Representative Western blots of 4E-BP1 phosphorylation (Thr37/46) are shown. Error bars represent standard error of the mean. **** = $p < 0.0001$, *** = $p < 0.001$, ** = $p < 0.01$, * = $p < 0.05$ by one-way ANOVA with Tukey correction in **(E)**, by unpaired Student's *t* test in **(B), (C), (D), (F), (G)**. *P*-value in **(H)** by Kolmogorov-Smirnov test.
